# Full-field strain distribution in hierarchical electrospun nanofibrous Poly-L(lactic) acid and Collagen based scaffolds for tendon and ligament tissue regeneration: a multiscale study

**DOI:** 10.1101/2023.06.01.543145

**Authors:** Alberto Sensini, Olga Stamati, Gregorio Marchiori, Nicola Sancisi, Carlo Gotti, Gianluca Giavaresi, Luca Cristofolini, Maria Letizia Focarete, Andrea Zucchelli, Gianluca Tozzi

## Abstract

Regeneration of injured tendons and ligaments (T/L) is a worldwide need. In this study electrospun hierarchical scaffolds made of a poly (L-lactic) acid and collagen blend were developed reproducing all the multiscale levels of aggregation of these tissues. Scanning electron microscopy, microCT and tensile mechanical tests were carried out, including a multiscale digital volume correlation analysis to measure the full-field strain distribution of electrospun structures. The principal tensile and compressive strains detected the pattern of strains caused by the nanofibers rearrangement, while the deviatoric strains revealed the related internal sliding of nanofibers and bundles. The results of this study confirmed the biomimicry of such electrospun hierarchical scaffolds, paving the way to further tissue engineering and clinical applications.

**Graphical Abstract:** 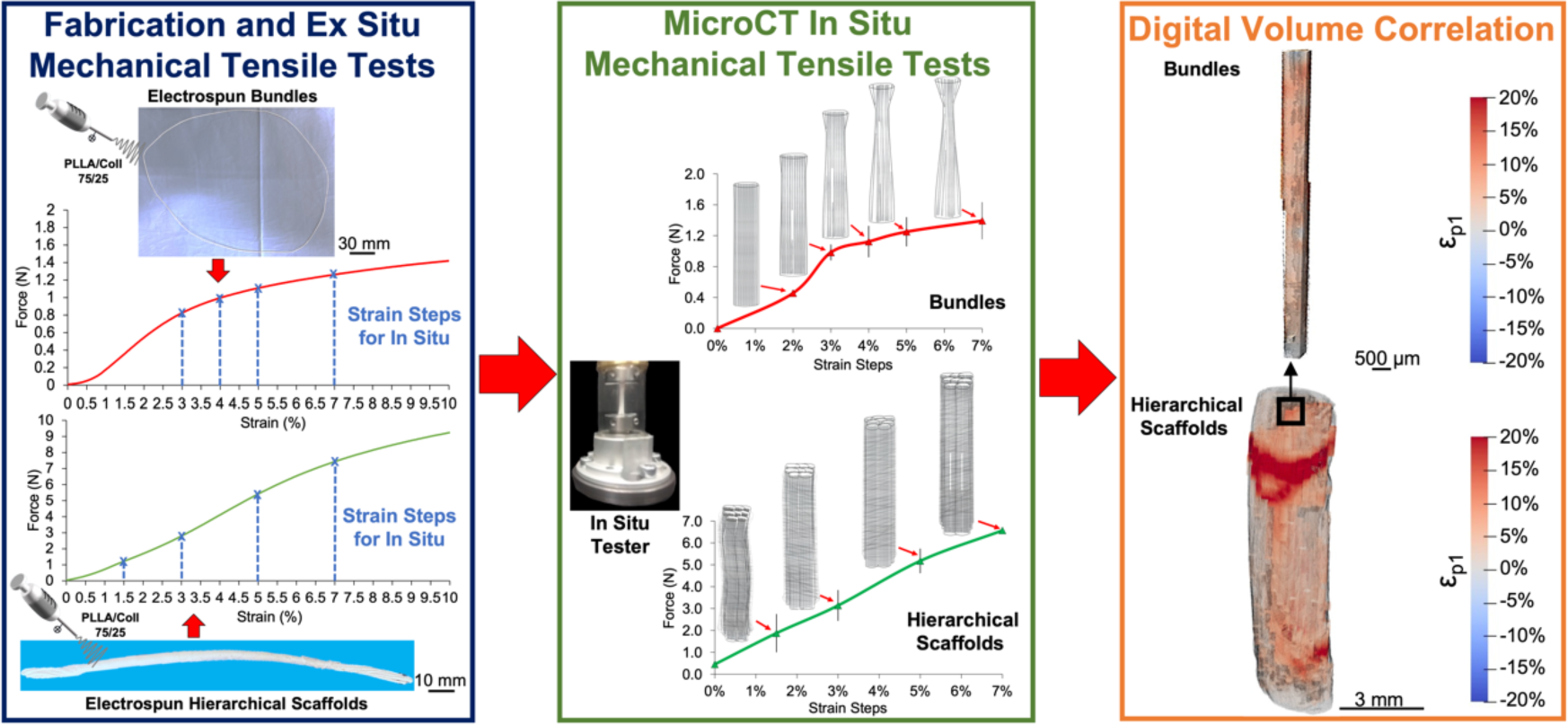

## **1.** Introduction

Injuries and ruptures of tendon/ligament (T/L) tissue represent one of the main challenges in modern orthopedics with approximately 30 million new injuries to these tissues worldwide and in constant increment each year [1]. The main cause of these lesions resides the high strains they are subjected to, that often damage their multiscale structure composed of nanometric fibrils of collagen type I, axially aligned and progressively aggregated in different hierarchical levels from the nano-up to the macroscale [2]. This complex morphology leads to non-linear mechanical properties, resulting from the interaction of these hierarchical levels [3]. To address the challenge of T/L regeneration, in the last twenty years tissue engineering has developed complex scaffolds to speed up their regeneration [4–6]. Among the various biofabrication techniques explored, electrospinning is for sure one of the most promising [7,8]. Sophisticated electrospun hierarchical structures, made of resorbable or biostable polymers, were developed to mimic T/L from the collagen fascicles [9–15], up to the whole tissue level [16–20], showing promising outcomes in enhancing cell proliferation and extracellular matrix (ECM) production by maintaining high morphological and mechanical biomimicry. Specifically, mechanical strains are a key aspect in the design of biomimetic scaffolds and it has been widely demonstrated how those contribute to guiding cells for the production of new ECM [21–23]. For this reason, several studies attempted to identify the two-dimensional strain patterns developed on the surface of natural or synthetic tissues mainly via digital image correlation (DIC) [24]. DIC investigations on T/L tissues were mostly focused on the human Iliotibial and Achilles tendons or on the Anterior Cruciate Ligament [25–30]. From the biofabrication side instead, DIC was used on electrospun mats for tissue engineering and enthesis (tendon to bone attachment) regeneration [31–34], successfully allowing to define their strain gradients. However, DIC is constrained to the measurement of superficial strains on the tested specimen and, to overcome this limitation, Digital Volume Correlation (DVC) was developed [35]. In brief, DVC relies on grayscale recognizable features, typically from X-ray micro computed tomography (microCT) images of materials subjected to progressive loading in situ, to measure volumetric full-field displacement and strain fields. The technique has been widely employed in musculoskeletal research [36]. Focusing on the T/L tissue instead, due to the high resolution and contrast required, microCT studies were performed mostly in static conditions by dehydrating samples, using contrast agents, or eventually performing phase-contrast synchrotron x-ray images [37–42]. To the authors’ knowledge, only one study has been carried out so far using DVC to study strain distributions in the rat enthesis [43]. Conversely, DVC analyses on electrospun materials and scaffolds are completely unexplored so far, due to the concomitant need of high resolution and the low X-ray absorption of polymeric fibrous materials. Thus, defining a DVC protocol to investigate the full-field strain distribution inside electrospun scaffolds is needed to finely tune their structure and mechanical properties, to optimally guide cells in their morphological/phenotype changes and in the production of new ECM during the early regeneration stages post-implantation.

Considering this background, the study aims at developing and applying the first microCT in situ protocol investigating the multiscale full-field strain distribution of electrospun structures via DVC. Results from single bundles and hierarchical scaffolds are also obtained and presented.

## 2. Materials and methods

### 2.1. Materials

Acid soluble collagen type I (Coll), extracted from bovine skin (Kensey Nash Corporation DSM Biomedical, Exton, USA) and Poly(L-lactic) acid (PLLA) (Lacea H.100-E, Mw = 8.4 × 104 g mol^−1^, PDI = 1.7, Mitsui Fine Chemicals, Dusseldorf, Germany) were used. 2,2,2-Trifluoroethanol (TFE), 1,1,1,3,3,3-Hexafluoro-2-propanol (HFIP) (Sigma-Aldrich, Staint Louis, USA) were used as solvents. For the crosslinking protocol N-(3-Dimethylaminopropyl)-N′-ethylcarbodiimide hydrochloride (EDC), N-Hydroxysuccinimide (NHS) and ethanol (Sigma-Aldrich, Staint Louis, USA) were used as received. The following polymeric blend solution was used: PLLA/Coll-75/25 (w/w) prepared from a 18% (w/v) solution of PLLA and Coll dissolved in TFE:HFIP = 50:50 (v/v).

### 2.2. Scaffolds Fabrication

To mimic the morphology of T/L fibrils and fascicles [2,44,45], PLLA/Coll-75/25 electrospun bundles of aligned nanofibers were produced as previously described [13,14,18]. To obtain ring bundles (RB) with a diameter in the range of human fascicle (500-650 μm), an industrial electrospinning machine (Spinbow srl, Bologna, Italy), equipped with a high-speed rotating drum collector (length = 405 mm, diameter = 150 mm; peripheral speed =19.6 m s^−1^; drum rotations = 2,500 rpm) and using an applied voltage of 22 kV, was used. To easily detach the nanofibers mats, the drum was covered with a sheet of polyethylene (PE) coated paper (Turconi S.p.A, Italy). The polymeric solution was delivered with four metallic needles (internal diameter = 0.51 mm, Hamilton, Romania), through PTFE tubes (Bola, Germany), using a feed rate of 0.5 mL h^−1^ controlled by a syringe pump (KD Scientific 200 series, IL, United States).

The needles-collector distance was of 200 mm; the sliding spinneret with the needles had an excursion of 180 mm, with a sliding speed of 1,500 mm min^−1^. The electrospinning session was set at 2 h at room temperature (RT) and with a relative humidity of 20–30%. After the electrospinning session, the mat was cut in circumferential stripes of 450 mm, wrapped up and pulled off the drum obtaining ring-shaped bundles (RB) of aligned nanofibers (Fig. 1B and 1C). To mimic the hierarchical structure of T/L [2,44,45] (Fig. 1A), hierarchical electrospun scaffold (EHS) (Fig. 1F) were assembled. Each RB was twisted in the middle and bent over itself (Fig. 1D). Then each assembly, composed of two folded RB, was covered with an electrospun epitenon/epiligament-like membrane, as previously described [17,18]. In brief, a second lab electrospinning machine (Spinbow srl, Bologna, Italy) equipped with a high-voltage power supply (FuG Elektronik GmbH, Schechen, Germany) and a syringe pump (KD Scientific Legato 100, Illinois, USA) was used to electrospin the solution. The two folded RB were placed in a custom-made setup, equipped with a flat plate aluminum collector, able to rotate the bundles during the electrospinning session. To produce the membrane RB were maintained in a static position and intermittently put in rotation (5 sessions of approximately 10 rpm for 30 sec every 20 min of stasis) (Fig. 1E). The PLLA/Coll-75/25 solution and the electrospinning parameters were the same as previously described. The scaffolds were finally crosslinked with a crosslinking solution of EDC and NHS 0.02 M in 95% ethanol, following a consolidated procedure [14].

**Figure 1.**
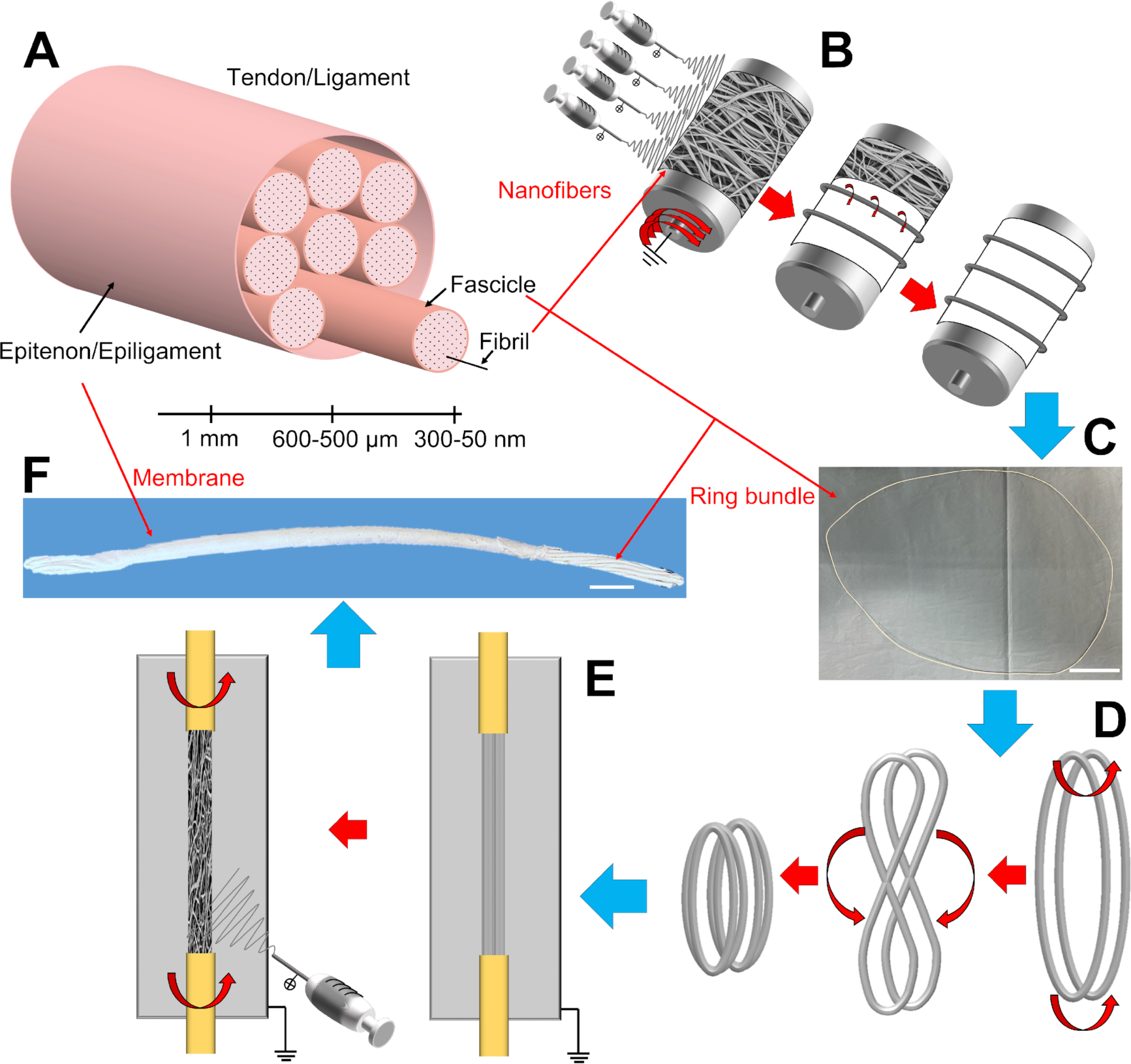
Electrospun scaffolds production. A) Hierarchical structure of tendons and ligaments. B) Electrospun ring bundles production. C) Example of a ring bundle (scale bar = 30 mm). D) Assembly of an EHS. E) Electrospun membrane production. F) Example of a final EHS obtained (scale bar = 5 mm).

### 2.3. SEM investigation

To visualize, at high-resolution, bundles and EHS surfaces, from the nano-up to the microscale, a Scanning Electron Microscopy (SEM) investigation, was carried out. Before the analysis, samples were gold-sputtered and then investigated with a SEM at 10 kV (SEM, Phenom Pro-X, PhenomWorld, Eindhoven, Netherlands). The opensource software ImageJ [46] was used to measure the diameters of 200 nanofibers (both for RB and EHS membranes; magnification = 8,000x) and the nanofiber diameter distribution was also computed. Both RB and EHS diameters were measured with an optical microscope (Axioskop, Zeiss, Pleasanton, CA, United States) equipped with a camera (AxioCam MRc, Zeiss, Pleasanton, CA, United States) and reported as mean and standard deviation of 20 measures. The nanofiber orientation was investigated with the Directionality plugin of ImageJ [47]. This approach allowed us to quantify the number of nanofibers within a given angle range from the axis, using a Local Gradient Orientation method, following a previously validated procedure [48]. For both bundles and EHS membrane, the analysis was performed on five images (magnification = 8,000x) along the scaffold’s axis and the results reported as mean and standard deviation between the five images.

### 2.4. Mechanical Characterization

To investigate the mechanical properties of the electrospun scaffolds and to set up the strain steps for the later in situ test, a mechanical tensile characterization of samples was carried out with a material testing machine (Mod. 4465, Instron, Norwood, United States) equipped with a ± 100 N load cell (Instron, Norwood, United States). The testing machine worked under displacement control to obtain a strain rate of 0.33% s^−1^. For each sample type (i.e., RB and EHS) dedicated capstan grips were used to reduce the stress concentration (see Figure 3). Before the test, each specimen was immersed in phosphate buffer saline (PBS). The mechanical performances of RB (n=5) and EHS (n=5) were tested using a monotonic ramp to failure (with a procedure adapted from the ASTM D1414 Standard) consistently with our previous study [49]. RB had a gauge length of 176 ± 1 mm while EHS had a gauge length of 90.0 ± 1 mm, caused by the specimen shrinkage after detachment from the drum collector and the later crosslinking. The force-displacement curves were converted to stress-strain graphs using two different approaches. In the first one, the apparent stress, calculated by dividing the force by the cross-sectional area of the specimen measured before the test, was plotted against strain. Whereas in the second one, the net stress was used to determine the mechanical properties of the specimen independently from its internal porosity. The net stress was calculated by dividing the apparent stress by the volume fraction (*v*) of the specimens. The volume fraction (*v*) was calculated by using the equation:

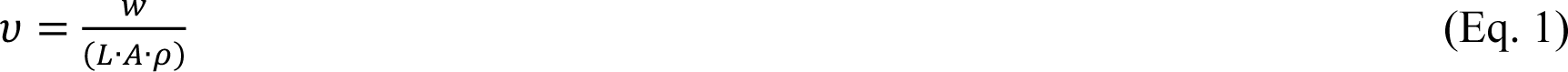

Where w is the weight of the specimen, L is length of the specimen, A is the cross-sectional area of the specimen, ρ is the density of the blend that, considering the density of PLLA (ρ_PLLA_ = 1.25 g cm^-3^) and of the Coll (ρ_Coll_ = 1.34 g cm^-3^), resulted in:

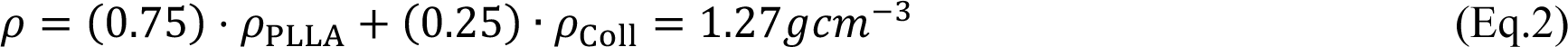

The weight of each specimen was calculated using a precision balance (AS 60/220.R2, Radwag, Pol). The following indicators were considered: Yield Force (F_Y_), Yield Stress (σ_Y_), Yield Strain (ε_Y_), Elastic Modulus (E), Failure Force (F_F_), Failure Stress (σ_F_), Failure Strain (ε_F_), Unit Work to Yield (W_Y_), Unit Work to Failure (W_F_). Moreover, by dividing by half the force of the RB, it was also possible to calculate F_Y_ and F_F_ of one of their branches, to set the strain values for the following in situ test on the single bundles (SB). To investigate the axial and transversal strains of the EHS membranes, 4K movies of the specimens, during the tensile test previously described, were acquired with a mobile phone camera (12-megapixel, Sony, JAP) synchronized with the testing machine. To allow an accurate measurement of the lengths of interest during the test, rulers were placed on each capstan grip (see Fig. 4E). Before the start of each tensile test, two zero-strain images were acquired as a reference. These images were used to calculate the axial (mean ± SD of 10 measures) and transversal (mean ± SD of 10 measures) length of the external membrane of the EHS under investigation. Then, after the mechanical elaboration of the curves, 5 high-resolution movie frames were selected corresponding to specific levels of strain (i.e. 1.5%, 3%, 5%, 7%, ε_Y_ and ε_F_) of the EHS previously calculated. For each image the axial and transversal length of the membrane were calculated with the same procedure reported for the zero-strain axial and transversal reference images. Finally, the axial (*ε_MA_*) and transversal (*ε_MT_*) strains of each membrane were calculated as follows:

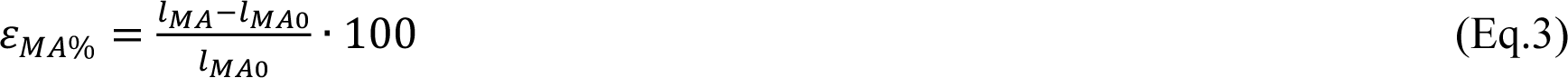

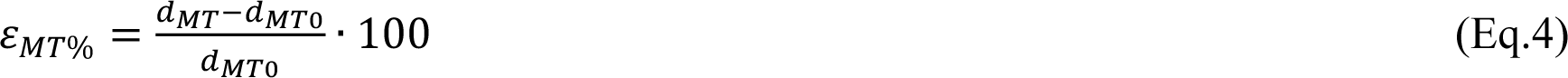

Where (*l_MA_)* is the mean axial length of the membrane corresponding to the investigated percentage of axial strain of the EHS specimen; (*l_MA0_)* is the mean axial length of the membrane at the zero-axial strain of the EHS specimen; (*d_MT_)* is the mean transversal diameter of the membrane at the investigated percentage of the axial strain of the EHS specimen; *(d_MT0_)* is the mean transversal diameter of the membrane at the zero-axial strain of the EHS specimen. All the measurements were taken with ImageJ.

### 2.5 MicroCT in situ protocol

Samples of SB (n=3) and EHS (n=3) underwent microCT in situ mechanical tests. The gauge length of the samples was measured by the in situ loading device, while the diameter by using ImageJ on optical microscope images. Each sample was immersed in PBS for 2 min (SB and EHS gauge length = 10 mm) before it was clamped in the microCT loading device (maximum actuator displacement = 5.5 mm; displacement rate = 0.001 mm s^-1^) (MTS in situ tester for Skyscan 1172, Bruker, Belgium). The first two consecutive microCT scans were acquired at the minimum strain allowed by the load cell sensitivity (0.45 Newton, corresponding to about 2% strain for SB and 0% strain for EHS) and were used to compute the DVC measurement uncertainties. Then, a series of progressive axial strain steps were imposed for SB (i.e. 3%, 4%, 5%, 7%) and for EHS (i.e. 1.5%, 3%, 5%, 7%). Strain levels were reached by imposing axial displacements based on the initial clamp-clamp distance, as measured on radiographic images on coronal and sagittal planes, while the tensile force was measured. At each strain step a 15 min stress-relaxation period was applied followed by a microCT acquisition [50]. SB samples were imaged with an applied voltage of 40 kV and current of 75 µA. The scan orbit was 180° with a rotation step of 0.8° and 4 frames averaged for each rotation angle, resulting in a voxel size of 13 µm and scan duration of ∼17 min. The scanning protocol for EHS samples was the same, except for increasing the voxel size to 9 µm to better discriminate between bundles and membrane. The image reconstruction was carried out with a modified Feldkamp algorithm by using the SkyScanTM NRecon software accelerated by GPU.

The region of interest (ROI) selection and morphometric analysis were all performed using SkyScan CT-Analyser software. The ROI extended 6 mm in length, which is 10 mm of gauge length minus 4 mm, to be away from steel clamps, avoiding metal artefacts. Membranes and SB could not be defined in a fully automatic way, by morphological operations, because of the tightening in traction and overlapping grey intensity distributions. For each sample and strain level, the following parameters were measured using the microCT post-processing software (CT-Analyser, Skyscan 1172, Bruker, Belgium):

1. Porosity *Po* (%): percentage of void space inside samples, was calculated by a 3D integrated Analysis (i.e., 3D morphometric parameters integrated for the whole volume) after defining a ROI that wraps the sample and binarizing the image with an automatic thresholding. For the EHS samples, a total (*Tot.Po*) and an internal (*Int.Po*) (i.e. excluding the membrane) porosity were defined (Fig. 2A);

**Figure 2.**
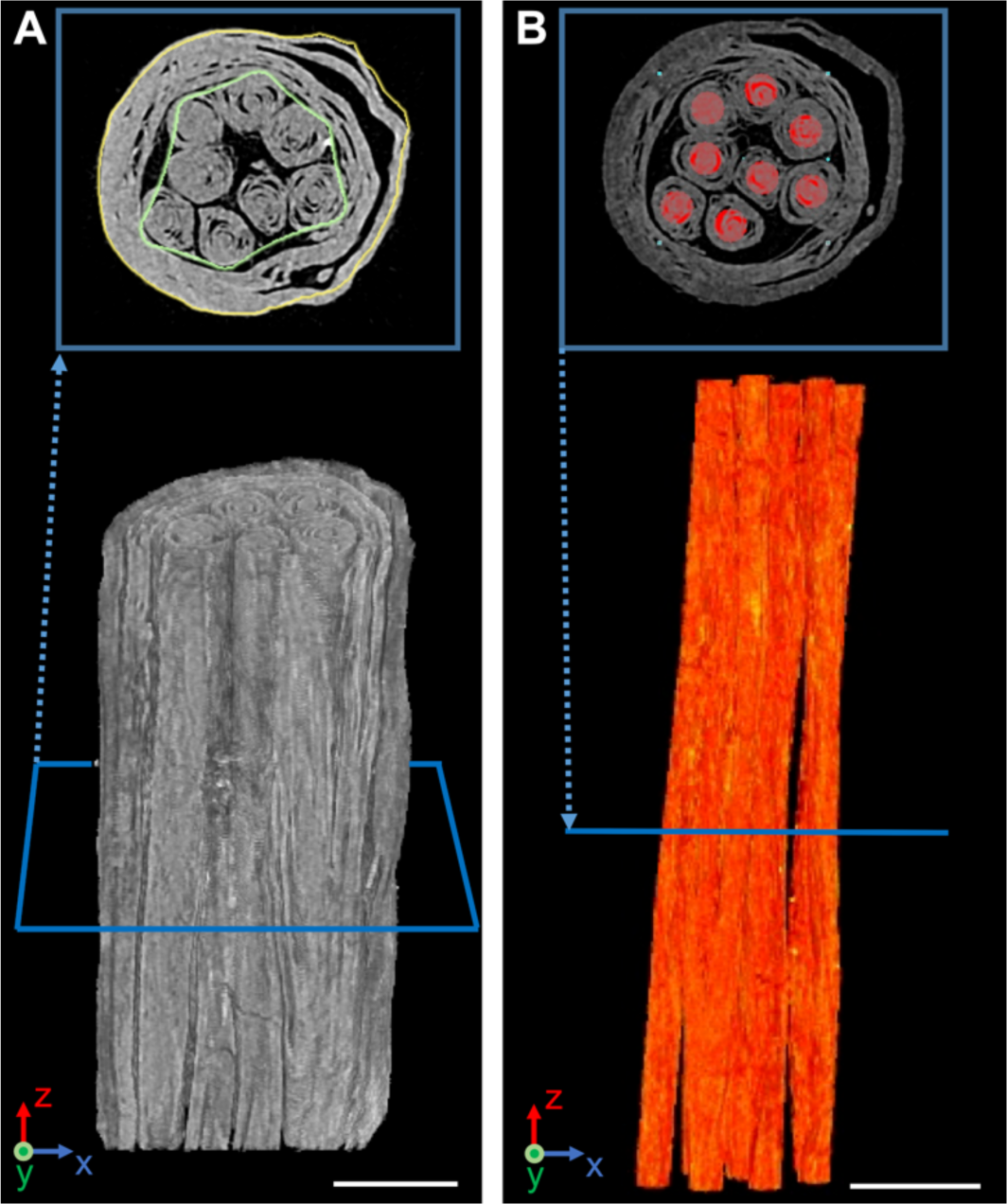
Workflow of the microCT morphometric analysis. A) EHS transversal section and rendering highlighting total area (contoured in yellow) and internal area (contoured in green); B) EHS transversal section and volume rendering highlighting objects (in red) running in bundles (scale bar = 1 mm).

2. MicroCT-computed cross-sectional area (i.e. *CT.Cr.Ar* in mm^2^): obtained by averaging the areas of the 450 transversal sections of the wrapping ROI described at point 1) net of porosity *Po* to consider resisting material only; SB orientation and tortuosity: calculated from morphological parameters of individual objects that follow the bundles’ major development in 3D space (Fig. 2B). Orientation *θ*(°) is defined as the object (i.e. bundle) angle respect to the loading (i.e. vertical axis Z in Fig. 2). It is 0° when parallel, 90° when orthogonal. Tortuosity (*τ*) instead, is defined as the object (i.e. bundle) equivalent length divided by the ROI vertical length (minimum value = 1, when the bundle is perfectly vertically aligned and without crimps) [51]. The same procedure was applied on the eight bundles inside each EHS.

Finally, thanks to the microCT-based morphometric parameters, the in situ strain-stress curves were calculated, considering the cross-sectional area of samples avoiding micro porosities visible from the microCT. In this way the microCT stress *CT*.σ (Ν) was defined as:

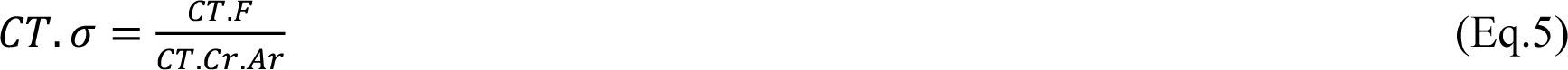

where *CT.F* (N) is the force recorded by the in situ tester during the experiment.

### 2.6. Digital Volume Correlation

Digital volume correlation was performed using the open-source software *spam* [52]. The correlation procedure in *spam* aims to measure a linear and homogeneous function, Φ, which is expressed in homogeneous coordinates and represented by a 4 × 4 deformation matrix that accounts for 3D affine transformations: translation, rotation, normal and shear strain. The formulation of the correlation algorithm is based on a gradient-based iterative procedure, which minimizes the difference between the reference and the deformed image, the latter being gradually corrected by a trial deformation function. The convergence criterion is based on the norm of the deformation function increment between two successive iteration steps, which was set here as: ‖δΦ‖ < 10^−4^. A maximum number of 500 iterations was also set as a limit to stop the iterative procedure in case that the convergence criterion was not satisfied.

A total DVC analysis was performed mapping the first scan (i.e., undeformed sample) with each of the remaining scans, as opposed to an incremental analysis which maps two consecutive load steps. To overcome the problem of the progressive large amounts of deformation for a total DVC analysis, for each pair of images an initial non-rigid registration was performed, which measured the overall average displacement, strain and rotation. This initial overall guess was then passed to a local approach, whereby independent cubic sub-volumes (i.e., correlation windows) were defined in the reference image (i.e., undeformed sample) and sought in the deformed image by applying the iterative procedure mentioned above. A Φ was computed in the centre of each window, yielding a field of deformation functions that mapped the reference to the deformed image. The size of the correlation windows and the number of measurement points depend on the texture of the imaged samples and define the spatial resolution of the measured field. For SB, a single run with a window size of 36 (i.e. 468 μm) pixels was enough to achieve a well-converged deformation field. For EHS samples, to achieve a good convergence in the local calculations, as well as a high spatially resolved deformation field a two-step approach of local DVC computations was performed with decreasing correlation window sizes from 100 (i.e. 900 μm) to 40 (i.e. 360 μm) pixels. In all cases an overlap of 50% was set between neighboring correlation sub-volumes.

Strains were obtained by extracting only the displacement part of the total fields of Φ and computing the transformation gradient tensor:

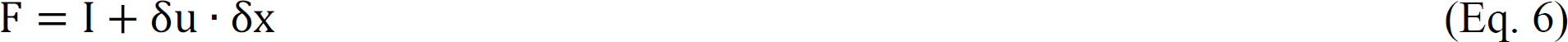

on Q8 shape functions linking 2 × 2 × 2 neighboring measurement points. Note that the displacement field was firstly smoothed by applying a 3D median filter of a 1 voxel radius. A polar decomposition of the transformation gradient tensor:

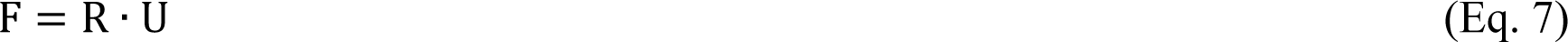

yielded the right stretch tensor U and the rotation tensor R for each Q8 element. The finite large-strain framework was used to calculate:

i) the principal strains ε_p1_ and ε_p3_ based on the diagonalization of the right Cauchy-Green deformation tensor:

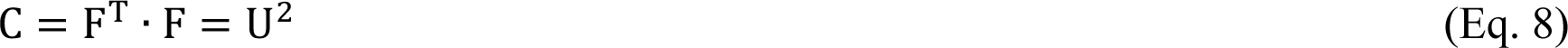

ii) the deviatoric strain ε_D_ based on a multiplicative decomposition of the stretch tensor U into a pseudo-isotropic and deviatoric part:

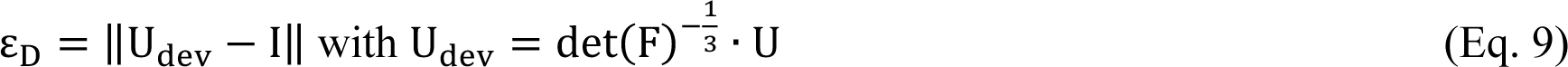

The level of uncertainty in the DVC procedure was estimated through a correlation analysis of the zero-strain scans [53]. As already mentioned, at the beginning of each test two scans of the undeformed sample were acquired. DVC was then run with the exact same parameters as the ones for the pair of images during the loading. The mean computed strain uncertainties were for SB: ε_p1_ = 0.2%, ε_p3_ = 0.2% and ε_D_ = 0.3%; for EHS instead: ε_p1_ = 0.3%, ε_p3_ = 0.3% and ε_D_ = 0.6%. DVC maps were overlaid onto microCT stacks using ImageJ and ParaView [54].

### 2.7. Statistical Analysis

The significance of differences between the apparent mechanical properties (i.e. forces) for the SB (n=5), RB (n = 5) and EHS (n = 5) was assessed with an ANOVA 1 unpaired parametric t-test with a Tukey post hoc (p>0.05, ns; p≤0.05, *; p≤0.01; **; p≤0.001, ***; p≤0.0001, ****). The significance of differences of apparent and net mechanical properties, between SB|RB (equal for single and ring bundles) and EHS was assessed with an unpaired parametric t-test with Welch’s correction. Instead, the comparison between the apparent and net mechanical properties of the same sample (i.e. SB|RB and HNES) was assessed with a ratio paired parametric t-test.

## 3. Results and Discussion

### 3.1. Morphology of bundles and EHS via SEM

Starting with a top-down approach, EHS (mean cross-sectional diameter = 2.7 ± 0.3 mm; mean length = 90 ± 1.0 mm; mean weight = 96 ± 10 mg) and bundles (mean cross-sectional diameter = 560 ± 93 μm; mean length = 176 ± 1.0 mm; mean weight = 35 ± 5 mg) had a similar morphology (Fig. 3A, 3C) and thickness of natural T/L reported in literature [2,3,44,45]. The SEM investigation revealed that PLLA/Coll-75/25 nanofibers of bundles (mean cross-sectional diameter = 0.238 ± 0.06 μm) (Fig. 2B) and membranes (mean cross-sectional diameter = 0.258 ± 0.08 μm) (Fig. 3D) were continuous, smooth and without defects such as beads. They also were in the same order of magnitude of T/L collagen fibrils [2,44,45]. The lower dispersion and diameters of the nanofibers of bundles compared with the ones of membranes, were consistent with the higher stretching caused by the drum collector (Fig. 3E). Interestingly, the Directionality analysis revealed a preferential axial orientation of the nanofibers of bundles and a slightly circumferential orientation of the membranes, consistently with our previous work [18]. The circumferential orientation of nanofibers in membranes confirmed the ability of the process to pack and tighten the structure. A relevant number of nanofibers of bundles were oriented in the range of 0°−12° (32.3% ± 2.2% of the total) from the bundle axis with a Gaussian-like distribution. A small number of nanofibers were oriented in the range of 81°−90° (6.0% ± 0.8% of the total). Conversely, the EHS membrane had lower number of nanofibers in the range of 0°−12° (6.8% ± 1.2% of the total) compared with the ones in the range of 81°−90° (16.1% ± 1.24% of the total). This analysis confirmed the morphological biomimicry of the bundles with the T/L fascicles and of the membranes with the epitenon/epiligament [2,43,44].

**Figure 3.**
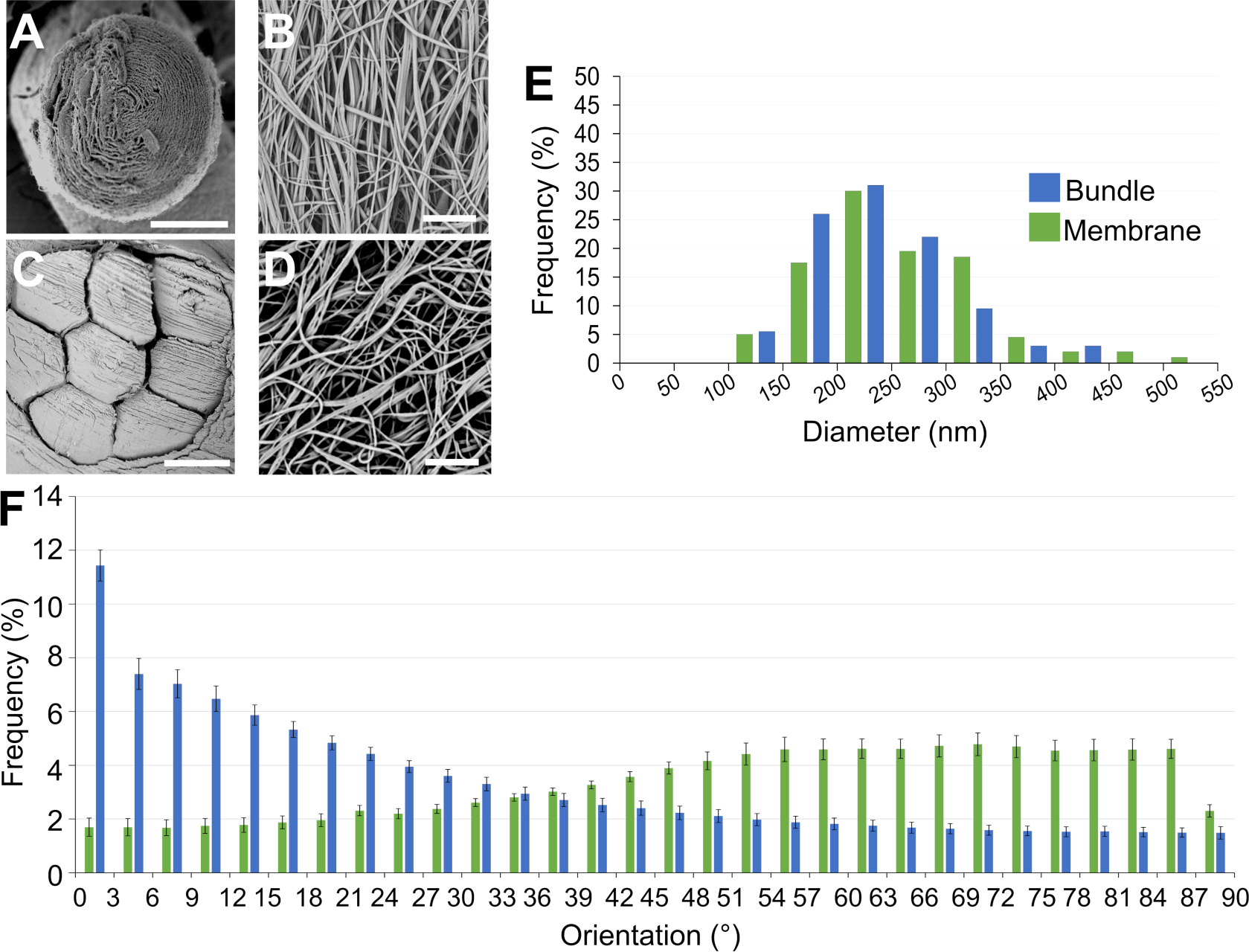
SEM images and morphological investigations of scaffolds. A) Bundle (scale bar = 300 μm; magnification = 500x). B) Nanofibers of the bundle (scale bar = 5 μm; magnification = 8000x). C) EHS (scale bar = 300 μm; magnification = 245x). D) Nanofibers of the membrane (scale bar = 5 m; magnification = 8000x). E) Nanofibers diameter distribution for the bundles and membranes. F) Orientation of the nanofibers of the bundles and EHS membranes. The Directionality histograms show the distribution of the nanofibers in the different directions. An angle of 0° means that the nanofibers were aligned with the axis of the scaffold, while an angle of 90° means that the nanofibers were perpendicular to the scaffold.

### 3.2 Mechanical properties of bundles and EHS

The mechanical properties of SB, RB and EHS, used to set the strain values for the in situ experiments, are reported in Fig. 4 and Table S2–S4.

**Figure 4.**
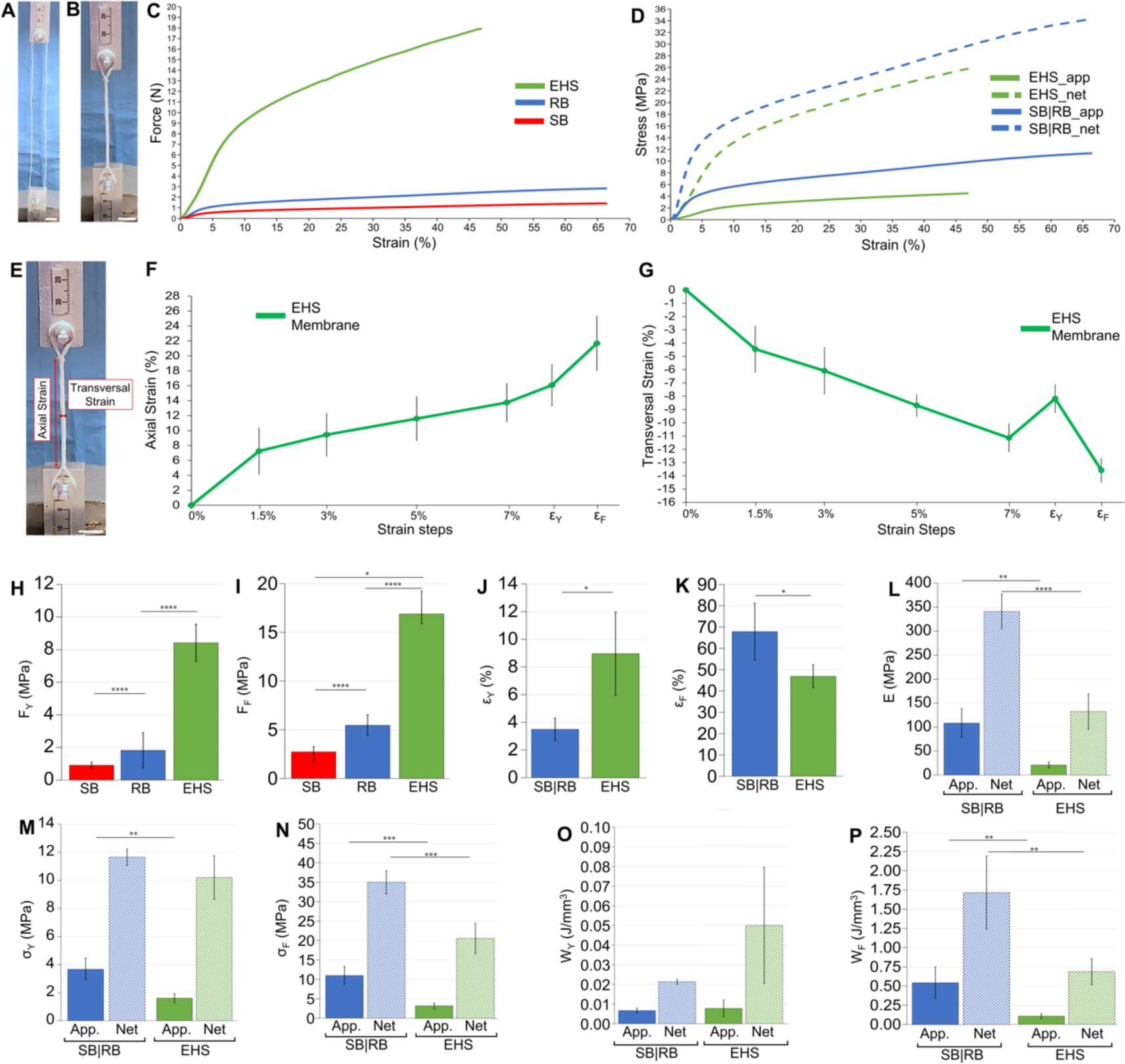
Mechanical tensile tests on bundles and EHS. A) Setup for testing RB (scale bar = 10 mm); B) setup for testing EHS (scale bar = 10 mm); C) typical force-strain curves for SB, RB and EHS; typical apparent and net stress-strain curves for SB|RB (same behavior being SB a branch of RB) and EHS; E) Example of axial and transversal strains (scale bar = 10 mm); F) mean and SD deviation of axial strain of EHS membranes at the different levels of strain of the in situ test including the yield and failure strain of EHS tensile tests; G) mean and SD deviation of transversal strain of EHS membranes at the different levels of strain of the in situ test including the yield and failure strain of EHS tensile tests; H) yield force of SB, RB and EHS (significance of differences of the ANOVA 1 showed with asterisks); I) failure force of SB, RB and EHS (significance of differences of the ANOVA 1 showed with asterisks); J) yield strain of SB|RB and EHS; K) failure strain of SB|RB and EHS; L) apparent and net elastic modulus for SB|RB and EHS; M) apparent and net yield stress for SB|RB and EHS; N) apparent and net failure stress for SB|RB and EHS; O) apparent and net work to yield for SB|RB and EHS; P) apparent and net work to failure for SB|RB and EHS. (H-N significance of differences from the unpaired parametric t-test with Welch’s correction reported with asterisks).

Both bundles and EHS showed a nonlinear toe region up to 2% strain, caused by the progressive stretching of the nanofibers under the applied load, followed by a linear elastic region, similar to the nonlinear behavior of fascicles and whole T/L [3,45]. Then, at different levels of strain (SB|RB: ε_Y_=3.5±0.11%; EHS: ε_Y_=8.97±3.01%), scaffolds showed a ductile region, partially caused by the bulk material properties and by the breakage of relevant number of nanofibers, up to their failure occurred at ε_F_ = 67.8±13.4% for SB/RB and at ε_F_ = 46.9±5.3% for EHS. The lower levels of failure strain for EHS were due to the folding procedure to obtain them. These failure strains were higher than those of natural fascicles and T/L [3,44], but can provide a safety factor in case of partial damage of scaffolds, with a relevant work absorption (Fig. 4P), preventing a premature implant failure. Other mechanical properties (Fig. 4 and Table S1), such as stress and elastic modulus, increased by approximately three times (for SB|RB) and six times (for EHS) passing from the apparent (SB|RB: σ_YApp._=3.67±0.80 MPa, σ_FApp._= 11.0±2.3 MPa, E_App._= 108±30 MPa, W_YApp._= 0.007±0.001 Jmm^-3^, W_FApp._= 0.54±0.20 Jmm^-3^; EHS: σ_YApp._= 1.61±0.31 MPa, σ_FApp._= 3.24±0.77 MPa, E_App._= 20.7±5.90 MPa, W_YApp._= 0.008±0.004 Jmm^-3^, W_FApp._= 0.11±0.03 Jmm^-3^) to the net ones (SB|RB: σ_YNet._=11.7±0.58 MPa, σ_FNet._= 35.0±3.0 MPa, E_Net._= 341±30 MPa, W_YNet._= 0.021±0.001 Jmm^-3^, W_FNet._= 1.72±0.50 J mm^-3^; EHS: σ_YNet._= 10.2±1.55 MPa, σ_FNet._= 20.6±3.87 MPa, E_Net._= 133±37 MPa, W_YNet._= 0.05±0.03 Jmm^-3^, W_FNet._= 0.69±0.17 Jmm^-3^) with statistically significant differences (Fig. 4 and Table S4). In fact, the net mechanical properties consider only the contribution of the volume fraction of the solid material that constitutes the scaffolds, without their internal porosities. Focusing on the net properties, all the mechanical values fell into the range of human fascicles [55] and whole T/L [3,45,56]. Moreover, except for a scalable increment in terms of forces with respect to SB|RB, the greater hierarchical complexity of EHS led to a decrement of stress, strain, elastic modulus and works with respect to bundles. This was caused by the increment of the internal adjustments of bundles with load and the higher porosity compared to SB|RB (i.e. for elastic modulus and stress and works). The epitenon/epiligament-inspired membranes of EHS successfully enclosed the internal bundles up to their failure (Fig. 4F, 4G and Table S5) showing a progressive increment of their axial strain at the different levels of EHS strain of the tensile test, in correspondence to the in situ strain steps. The transversal strains instead, showed a progressive reduction, caused by the striction and adjustments of the internal bundles up to 7% of EHS strain, then increasing before EHS ε_Y_ and finally reducing again to EHS ε_F_. The increment in transversal strain in the range 7% - ε_Y_ was due to the tendency of the internal bundles to follow the capstan grips external diameter.

### 3.3 Morphology and mechanics of bundles and EHS via microCT in situ tests

Values of strains, loads and morphological parameters from the microCT *in situ* test are shown in Fig. 5 and listed in Table S6.

**Figure 5.**
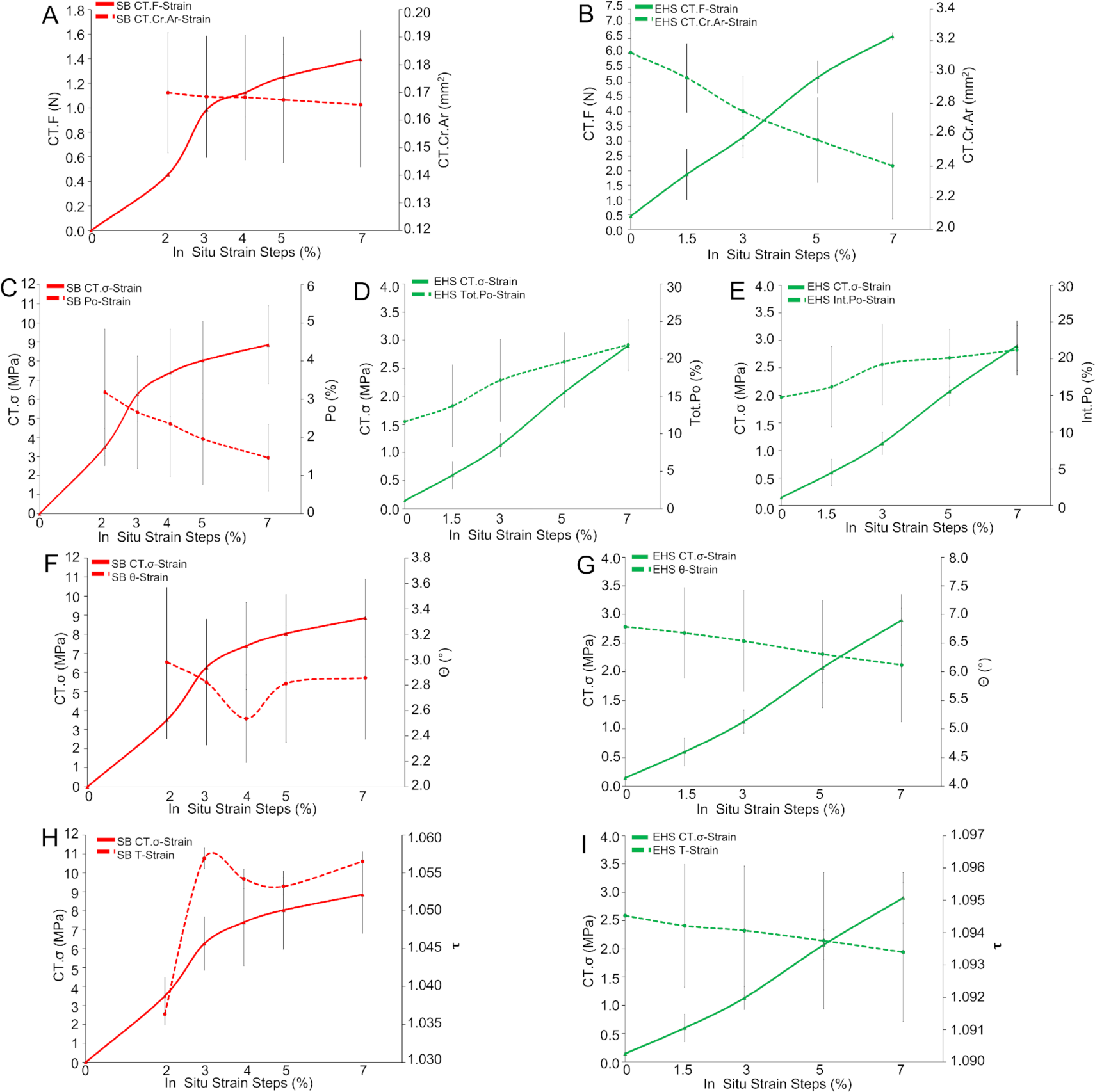
Evolution (mean and SD at the different in situ strain steps) of SB (red lines) and EHS (green lines) morphometric parameters and mechanical characteristics in comparison with the corresponding *CT.F* and *CT.σ*: A) SB *CT.F* and *CT.Cr.Ar*; B) EHS *CT.F* and *CT.Cr.Ar*; C) SB *CT.σ* and *Po*; D) EHS *CT.σ* and *Tot.Po*; E) EHS *CT.σ* and *Int.Po*; F) SB *CT.σ* and *θ*; G) EHS *CT.σ* and *θ*; H) SB *CT.σ* and *τ*; I) EHS *CT.σ* and *τ*.

Load-strain curves (*CT.F* in Fig.5A, 5B) were consistent with those of the reference (ex situ) mechanical characterization (Fig. 4C) both for SB and EHS, supporting the validity of the microCT *in situ* protocol. The measurement of the cross-sectional area of the material during the in situ tensile steps and the net of microporosity, at each strain level, highlighted the expected striction phenomenon (*CT.Cr.Ar* in Fig. 5A, 5B), allowing also to follow closely the evolution of stress (*CT.σ* in Fig. 5C-I). This revealed that, during microCT acquisition, the SB mechanical response was almost in the yielding region, while EHS was still in the elastic one. The cross-sectional area restriction corresponded to a decrement in microporosity only for SB (intra-bundle voids, *Po* in Fig. 5C) and to an increase for EHS (intra-bundle and inter-bundles voids, *Int.Po* in Fig. 5E), due to the reduction of tortuosity and the parallel increament in separation between bundles with increasing strain. SB tortuosity (*τ* in Fig. 5H) and orientation (*θ* in Fig. 5F) showed no trend with strain and lower average-on-strains values (1.05 and 2°, respectively) with respect to EHS, in which instead they slightly decreased with strain (Fig. 5G, 5I). This can be related to a stretched structural arrangement in SB, that is in fact yielding, while to a progressive alignment on loading direction in EHS, that is still elastically deforming.

### 3.4 Digital Volume Correlation Analysis

The DVC successfully measured, starting from the displacement fields, the full-field strain distribution both on SB and EHS (Table 1, Table S7, Table S8, Fig. 7 and Fig. 8). The uncertainties calculated where approximately one order of magnitude lower of the mean strain at the yielding point of both SB and EHS. These values are consistent for the strain analysis of such viscoelastic materials.

**Table 1.** Mean ± SD of axial displacements and DVC ε_p1_, ε_p3_, ε_D_ strains between the tested samples of the same category, together with their maximum and minimum values.

| Strain Steps<br>(SB) | 2-3% | 2-4% | 2-5% | 2-7% |
| --- | --- | --- | --- | --- |
| Disp (mm) | 0.08±0.03 | 0.14±0.04 | 0.21±0.08 | 0.35±0.12 |
| Disp<sub>max</sub> (mm) | 0.16±0.07 | 0.25±0.06 | 0.35±0.12 | 0.60±0.12 |
| $\epsilon_{p1}$ (%) | 1.62±0.30 | 2.20±0.13 | 3.05±0.20 | 4.78±0.56 |
| $\epsilon_{p1max}$ (%) | 6.91±2.28 | 8.43±1.63 | 11.20±4.12 | 14.57±6.89 |
| $\epsilon_{p3}$ (%) | -0.97±0.30 | -1.01±0.19 | -1.36±0.12 | -1.85±0.05 |
| $\epsilon_{p3min}$ (%) | -4.28±0.97 | -4.50±2.33 | -5.24±1.28 | -6.11±1.56 |
| $\epsilon_D$ (%) | 1.92±0.46 | 2.4±0.26 | 3.28±0.18 | 4.86±0.58 |
| $\epsilon_{Dmax}$ (%) | 5.13±1.48 | 6.92±0.94 | 9.76±2.69 | 14.12±3.82 |
| **Strain Steps**<br>**(EHS)** | **0-1.5%** | **0-3%** | **0-5%** | **0-7%** |
| Disp (mm) | 0.06±0.03 | 0.13±0.05 | 0.23±0.06 | 0.38±0.06 |
| Disp<sub>max</sub> (mm) | 0.12±0.04 | 0.22±0.09 | 0.36±0.09 | 0.58±0.17 |
| $\epsilon_{p1}$ (%) | 2.19±1.23 | 3.34±1.20 | 5.15±1.38 | 7.26±0.75 |
| $\epsilon_{p1max}$ (%) | 10.98±3.41 | 22.74±5.83 | 30.54±5.67 | 40.17±3.31 |
| $\epsilon_{p3}$ (%) | -2.41±0.56 | -4.10±0.64 | -6.27±1.63 | -8.35±3.52 |
| $\epsilon_{p3min}$ (%) | -13.06±1.69 | -18.50±2.86 | -24.90±5.19 | -27.00±6.04 |
| $\epsilon_D$ (%) | 3.35±1.23 | 5.45±0.34 | 8.38±0.80 | 10.75±1.30 |
| $\epsilon_{Dmax}$ (%) | 15.22±3.99 | 22.80±6.33 | 32.40±3.88 | 37.77±4.33 |

A hypothesis of the rearrangement of SB internal nanofibers during the in situ test can be summarized as follows (see Fig. 6A and Fig. 7).

**Figure 6.**
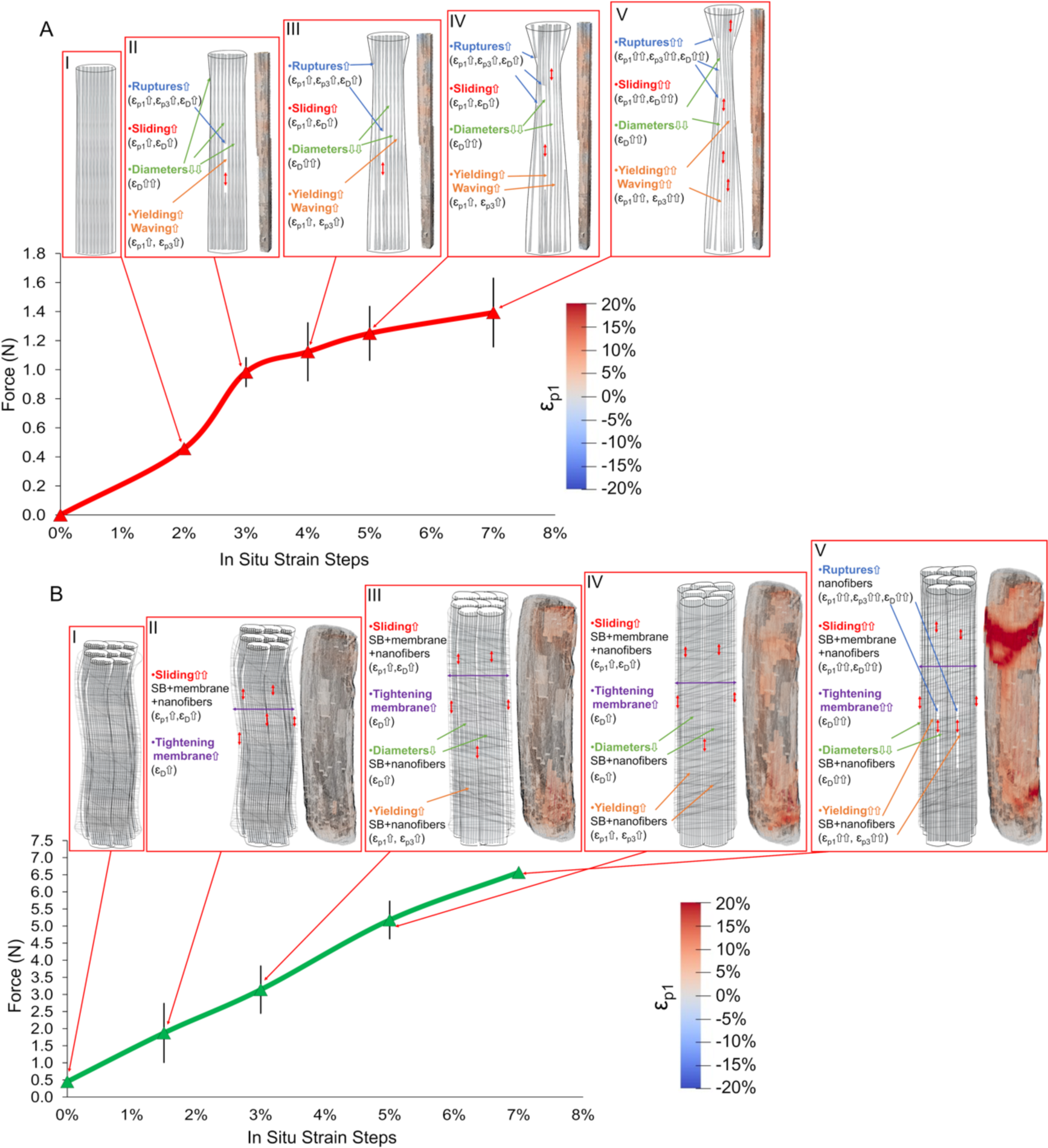
DVC strain evolution with nanofibers and scaffolds organization during the different in situ mechanical test. A) Strain evolution of SB at the different strain steps: AI) 2%; AII) 3%; AIII) 4%; AIV) 5%; AV) 7%. B) Strain evolution of EHS at the different strain steps: BI) 0%; AII) 1.5%; AIII) 3%; AIV) 5%; AV) 7%.

In the first step the SB are already in the linear region where, having passed the nonlinear toe region, nanofibers are a bit stretched and aligned (Fig. 6AI). At 3% the yielding point is reached, and nanofibers, and the wrapped mat layers that compose each SB, rise their stretching and ε_p1_ (Fig. 6AII). The first ruptures of nanofibers and layers occur, enhancing ε_D_, while nanofibers progressively reduce their diameters, showing some relaxation. These phenomena increase the compressive regions and ε_p3_ also causing an amplification of sliding (ε_D_) in the subsequent strain steps, progressively amplifying the phenomena previously described (Fig. 6AIII-6AV).

Specifically, the SB (Table 1, Fig. 7 and Table S7) showed, as expected, increasing ε_p1_ strains during the in situ test reaching local values of ε_p1_ = 4.78 ± 0.56% at 7% step, but with maximum peak of ε_p1max_ = 14.57±6.98% (from 10 up to 3 times higher strain compared to the apparent strain values) (Fig. 7B). Consistently, ε_p3_ confirmed a progressive striction of the cross-section of SB with the elongation/yielding of the internal nanofibers with mean negative values of ε_p3_ = -1.85±0.05% (ε_p3min_ = - 6.11±1.56%) at 7% step (Fig.7C). However locally, SB also exhibited positive values of ε_p3_ that could be probably caused by the concomitant presence of: i) internal reorganization of the nanofibers and layers of the original electrospun mat used for generating the bundles; ii) local relaxation of groups of nanofibers as they yield. Similarly, the deviatoric strain ε_D_ confirmed a progressive sliding of the internal electrospun layers of the wrapped mat of bundles, their nanofibers and the evolution of the internal porosities of SB, reaching mean values of ε_D_ = 4.86±0.58% (Fig. 7D), with local maxima ε_Dmax_ = 14.12±3.82%. Considering the PLLA/Coll nanofibers, these were not resolvable at the voxel size achieved from microCT (i.e. SB = 13 μm; EHS = 9 μm). However, their overall rearrangement inside the SB volume can explain the strain behavior of SB, which is also supported by the morphometric investigation (see Fig. 5 and Table S6).

**Figure 7.**
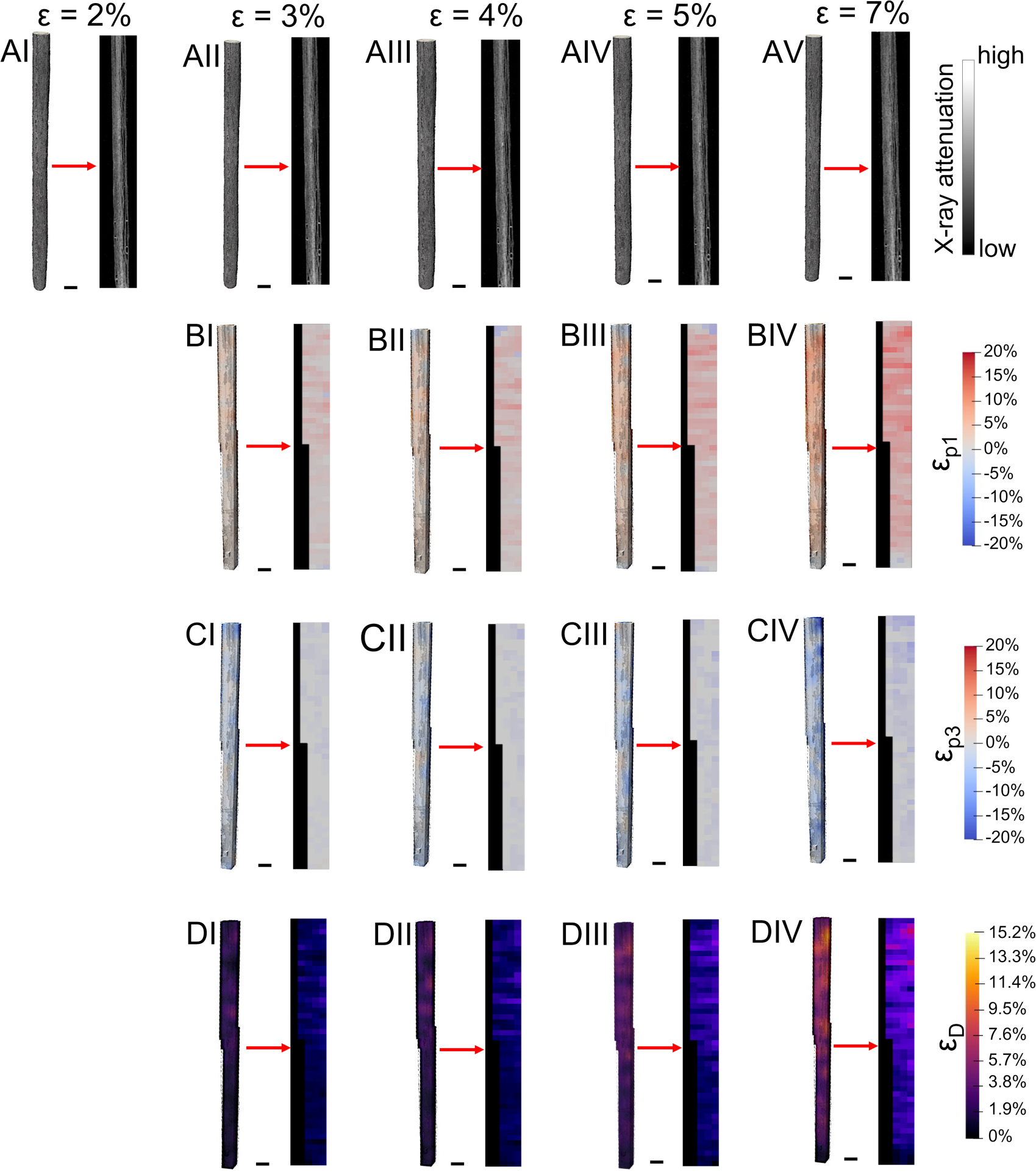
Evolution during the in situ test of the DVC strain fields for a representative SB 3D volume and its internal cross-section (scale bar = 500 μm): A) reconstructed 3D volume renderings and central vertical cross-section for the different strain steps; B) 3D volume renderings and central vertical cross-section for the different strain steps of ε_p1_; C) 3D volume renderings and central vertical cross-section for the different strain steps of ε_p3_; D) 3D volume renderings and central vertical cross-section for the different strain steps of ε_D_.

For EHS instead, the strain evolution must consider additional phenomena (Fig.6B and Fig.8). At the zero-strain in this case, the nanofibers and bundles are waved, being in the toe region of scaffolds (this happens because the initial pre-load is distributed in the internal bundles), while the nanofibers in the membrane are partially circumferentially oriented (Fig.6BI). In the second step, nanofibers and bundles progressively increase their alignment producing some sliding and tightening of the membrane and causing an increment of ε_D_ and ε_p1_ (Fig. 6BII). In the third step, nanofibers and bundles are now aligned experiencing an incremental stretching/sliding of the internal bundles and nanofibers, and with a parallel tightening of the membrane (with a rise of ε_D_ and ε_p1_) (Fig. 6BIII). These phenomena will also cause an overall reduction of EHS diameter (increment of ε_D_) but also some yielding of the smallest nanofibers (ε_p1_ and ε_p3_ up). In the last steps, being EHS close to yielding, all the previous phenomena are progressively amplified including the rupture of bunches of nanofibers (raising all the strains) (Fig. 6BIV and 6BV).

**Figure 8.**
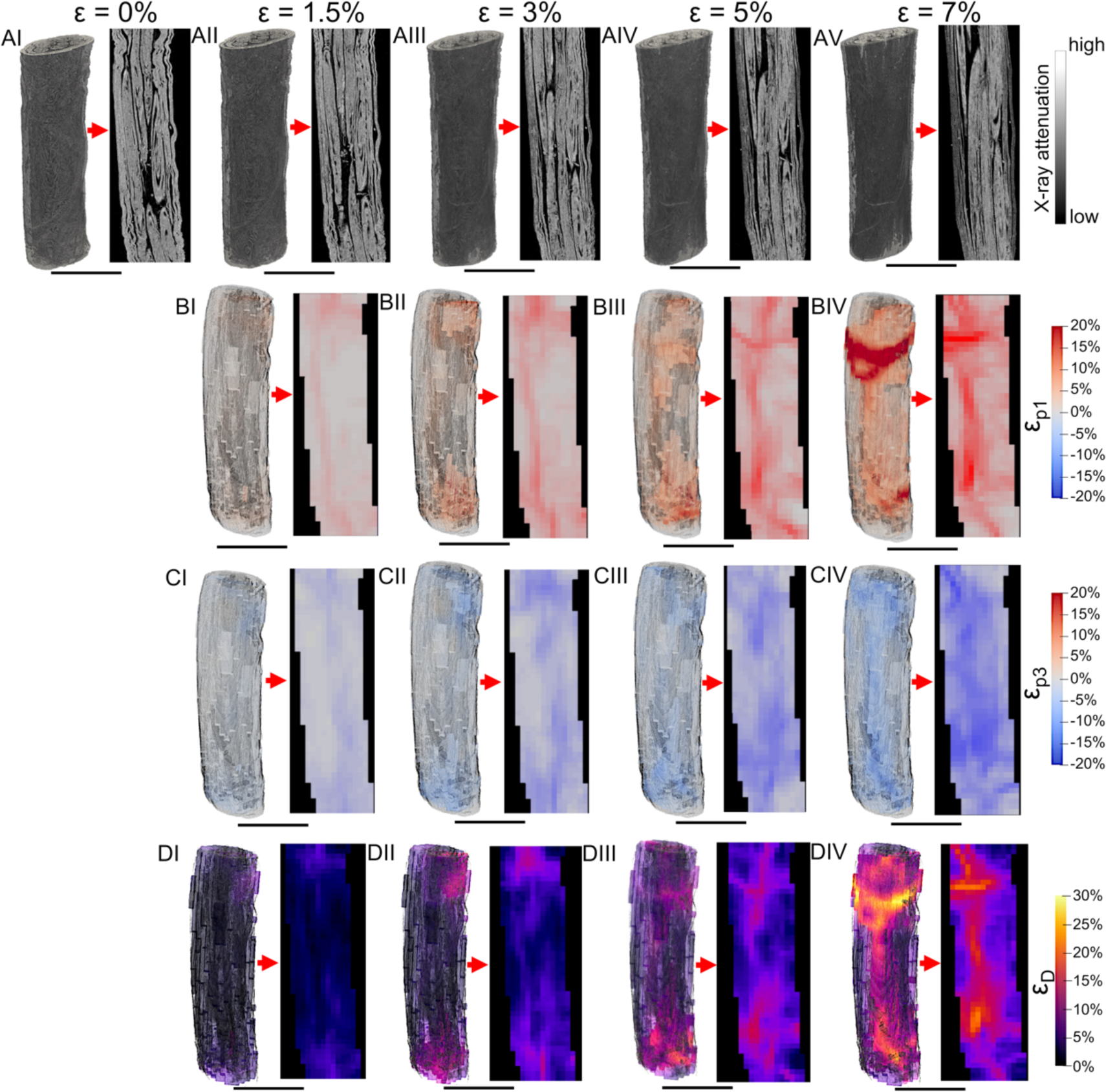
Evolution during the in situ test of the DVC strain fields for a representative EHS 3D volume and its internal cross-section (scale bar = 3 mm): A) reconstructed 3D volume renderings and central vertical cross-section for the different strain steps; B) 3D volume renderings and central vertical cross-section for the different strain steps of χ_p1_; C) 3D volume renderings and central vertical cross-section for the different strain steps of ε_p3_; D) 3D volume renderings and central vertical cross-section for the different strain steps of ε_D_.

More specifically, in EHS (Fig. 8 and Table 1 and Table S8), the strain-guided evolution of the internal bundles and void spaces were detected from the images. The ε_p1_ showed a progressive increment of mean values up to ε_p1_ = 7.26±0.75% (ε_p1max_ = 40.17±3.31%) at 7% of apparent strain. EHS also showed compressive regions with a mean ε_p3_ = - 8.35±3.52% at 7% step (ε_p3min_ = -27.00±6.04%) (Fig. 8B). Conversely in the regions were ε_p3_ exhibits negative values, it is possible to see negative values of ε_p1_, suggesting a similar progressive damage/local relaxation and reorganization of the internal bundles and nanofibers of the membrane. Moreover, due to their reorganization during the different strain steps, the preferential transversal alignment of the nanofibers of the membrane contributed to a progressive tightening of the EHS, increasing the negative values of ε_p3_. These data are also in accordance with the morphometric increment of internal porosity and θ of EHS, the progressive decrement of (τ) (Fig. 5 and Table S6) and the slightly increment of the transversal strain of the membranes between ε_Y_ and the 7% step (Fig. 4G). All these adjustments of EHS internal bundles were also confirmed by the mean values of ε_D_ = 10.75±1.30% (Fig. 8D) in correspondence of their maximum sliding (ε_Dmax_ = 10.75±1.30%).

The DVC full-field strain distribution of SB and EHS produced a strain behavior similar to that experienced in the natural T/L tissue counterpart by using DVC [43], DIC [26,28,57,58] and finite element models [59]. The progressive stretching and reorganization of internal collagen fibrils/fascicles of T/L during physiological activities is responsible for the nonlinear behavior of their stress/strain curves. This characteristic is typically visible using DIC on T/L and resulting into inhomogeneous strain patterns that follow the local stretch/relaxation during tensile test [26,28,57,58]. Moreover, it is also well established that the collagen fibrils in T/L start damaging in the linear region of the stress/strain curve in a specific point defined as “inflection point” [60]. This point identifies the region where the curve transits from stiffening to softening. This behavior of internal yield/relaxation in both SB and EHS from the current study is therefore consistent with natural T/L and their fascicles.

The measured full-field mechanics of the examined scaffolds contributed to better explain the morphological changes and elongation of fibroblasts and tenocytes previously detected during static [13,14] and dynamic cultures in bioreactor [18]. These data contribute to explain the reason why these scaffolds can interact this cells guiding their morphological changes in shape and orientation [18].

This study confirmed the promising morphological and mechanical performance of such scaffolds for T/L tissue engineering, also showing some limitations mostly related to the resolution of the microCT scans and the clamping setup. In fact, as mentioned, the resolution of the microCT system used in this study did not allow detection of individual nanofibers, as well as their internal micro/nano porosities much smaller than the nominal voxel size (scans voxel size = 9-13 μm). Also, the step-wise nature of the in situ test (i.e. 30 minutes for each strain step) contributed to partial relaxation inside the scaffolds due to the intrinsic viscoelasticity of these polymeric materials. All samples of SB and EHS showed peaks of strain in close proximity of the clamps of the in situ loading device confirming, as expected, strain concentrations due to the clamping setup. Future studies will require reduced scanning times coupled with higher resolutions, for example using synchrotron x-ray computed tomography, to visualize and measure phenomena at the nanofiber and using dedicated capstan grips to minimize strain concentration.

## 4. Conclusion

In this study, biomimetic PLLA/Coll-based electrospun scaffolds for T/L tissue engineering were successfully produced and characterized with techniques such as in situ microCT and DVC allowing, for the first time, to measure the 3D full-field strain distribution inside electrospun materials. The scaffolds mimicked the multiscale morphological and mechanical behavior of the natural collagen fibril/fascicles to the whole T/L tissue. The findings of this study will provide fundamental insights for future research on electrospinning and regenerative medicine, to better understand the complex interplay between nanofibrous structure/mechanics and how this can optimally drive cell fate in vivo.

## 5. CRediT Authorship Contribution Statement

**Alberto Sensini**: Conceptualization, Electrospun scaffolds production, Mechanical tests, SEM morphological investigations, Methodology for the microCT in situ tests and formal analysis, Help in DVC analysis, Writing – review & editing, Funding acquisition. **Olga Stamati**: DVC analysis and microCT images rendering, Writing – review & editing. **Gregorio Marchiori**: Methodology for the microCT in situ tests and formal analysis, MicroCT morphological investigations, Writing – review & editing. **Nicola Sancisi**: Methodology for the microCT in situ tests and formal analysis, Writing –review & editing. **Carlo Gotti**: Mechanical tests, Writing – review & editing. **Gainluca Giavaresi:** Supervision of the microCT facility, Writing – review & editing. **Maria Letizia Focarete**: Writing – review & editing, Funding acquisition. **Luca Cristofolini**: Writing – review & editing, Funding acquisition. **Andrea Zucchelli**: Conceptualization, Writing – review & editing, Funding acquisition. **Gianluca Tozzi**: Conceptualization, Methodology for the microCT in situ tests, Supervision of the DVC analysis, Writing – review & editing.

## Acknowledgments

The study was kindly supported by the University of Bologna Proof Of Concept Grant and partially the Horizon Europe Marie Skłodowska-Curie Postdoctoral Fellowship (Grant No. 101061826 3NTHESES project https://cordis.europa.eu/project/id/101061826). Type I collagen was kindly provided by Kensey Nash Corporation d/b/a DSM Biomedical (Exton, USA). Edward Andò is acknowledged for the useful suggestions in the use of *spam* software. Ilaria Fuoco and Gaia Prezioso are acknowledged for the help in samples preparation, in the experimental part and data analysis.

## 7. Data Availability

Data will be made available on request.

## References

[1] Abbah SA, Spanoudes K, O’Brien T, Pandit A, Zeugolis D. Assessment of stem cell carriers for tendon tissue engineering in pre-clinical models. Stem Cell Res Ther 2014;5:1–9. doi:10.1186/scrt426.

[2] Kastelic J, Galeski A, Baer E. The Multicomposite Structure of Tendon. Connect Tissue Res 1978;6:11–23. doi:10.3109/03008207809152283.

[3] Goh KL, Listrat A, Béchet D. Hierarchical mechanics of connective tissues: Integrating insights from nano to macroscopic studies. J Biomed Nanotechnol 2014;10:2464–507. doi:10.1166/jbn.2014.1960.

[4] Shiroud Heidari B, Ruan R, Vahabli E, Chen P, De-Juan-Pardo EM, Zheng M, et al. Natural, synthetic and commercially-available biopolymers used to regenerate tendons and ligaments. Bioact Mater 2023;19:179–97. doi:10.1016/j.bioactmat.2022.04.003.

[5] Rinoldi C, Kijeńska-Gawrońska E, Khademhosseini A, Tamayol A, Swieszkowski W. Fibrous Systems as Potential Solutions for Tendon and Ligament Repair, Healing, and Regeneration. Adv Healthc Mater 2021;10. doi:10.1002/adhm.202001305.

[6] No YJ, Castilho M, Ramaswamy Y, Zreiqat H. Role of Biomaterials and Controlled Architecture on Tendon/Ligament Repair and Regeneration. Adv Mater 2020;32:1–16. doi:10.1002/adma.201904511.

[7] Sensini A, Cristofolini L. Biofabrication of Electrospun Scaffolds for the Regeneration of Tendons and Ligaments. Materials (Basel) 2018;11:1963. doi:10.3390/ma11101963.

[8] Sensini A, Massafra G, Gotti C, Zucchelli A, Cristofolini L. Tissue Engineering for the Insertions of Tendons and Ligaments: An Overview of Electrospun Biomaterials and Structures. Front Bioeng Biotechnol 2021;9:1–23. doi:10.3389/fbioe.2021.645544.

[9] Brennan DA, Conte AA, Kanski G, Turkula S, Hu X, Kleiner MT, et al. Mechanical Considerations for Electrospun Nanofibers in Tendon and Ligament Repair. Adv Healthc Mater 2018;1701277:1–31. doi:10.1002/adhm.201701277.

[10] Bosworth LA, Alam N, Wong JK, Downes S. Investigation of 2D and 3D electrospun scaffolds intended for tendon repair. J Mater Sci Mater Med 2013;24:1605–14. doi:10.1007/s10856-013-4911-8.

[11] Bosworth LA, Rathbone SR, Bradley RS, Cartmell SH. Dynamic loading of electrospun yarns guides mesenchymal stem cells towards a tendon lineage. J Mech Behav Biomed Mater 2014;39:175–83. doi:10.1016/j.jmbbm.2014.07.009.

[12] Pauly HM, Kelly DJ, Popat KC, Trujillo NA, Dunne NJ, McCarthy HO, et al. Mechanical properties and cellular response of novel electrospun nanofibers for ligament tissue engineering: Effects of orientation and geometry. J Mech Behav Biomed Mater 2016;61:258–70. doi:10.1016/j.jmbbm.2016.03.022.

[13] Sensini A, Gualandi C, Cristofolini L, Tozzi G, Dicarlo M, Teti G, et al. Biofabrication of bundles of poly(lactic acid)-collagen blends mimicking the fascicles of the human Achille tendon. Biofabrication 2017;9. doi:10.1088/1758-5090/aa6204.

[14] Sensini A, Gualandi C, Zucchelli A, Boyle L, Kao AP, Reilly GC, et al. Tendon Fascicle-Inspired Nanofibrous Scaffold of Polylactic acid/Collagen with Enhanced 3D-Structure and Biomechanical Properties. Sci Rep 2018;8:1–15. doi:https://doi.org/10.1038/s41598-018-35536-8.

[15] Sensini A, Santare MH, Eichenlaub E, Bloom E, Gotti C, Zucchelli A, et al. Tuning the Structure of Nylon 6,6 Electrospun Bundles to Mimic the Mechanical Performance of Tendon Fascicles. Front Bioeng Biotechnol 2021;9:1–12. doi:10.3389/fbioe.2021.626433.

[16] Laranjeira M, Domingues RMA, Costa-Almeida R, Reis RL, Gomes ME. 3D Mimicry of Native-Tissue-Fiber Architecture Guides Tendon-Derived Cells and Adipose Stem Cells into Artificial Tendon Constructs. Small 2017;13:1–13. doi:10.1002/smll.201700689.

[17] Sensini A, Gualandi C, Focarete ML, Belcari J, Zucchelli A, Boyle L, et al. Multiscale hierarchical bioresorbable scaffolds for the regeneration of tendons and ligaments. Biofabrication 2019;11:35026. doi:10.1088/1758-5090/ab20ad.

[18] Sensini A, Cristofolini L, Zucchelli A, Focarete ML, Gualandi C, de Mori A, et al. Hierarchical electrospun tendon-ligament bioinspired scaffolds induce changes in fibroblasts morphology under static and dynamic conditions. J Microsc 2020;277:160–9. doi:10.1111/jmi.12827.

[19] Xie X, Xu J, Lin J, Jiang J, Huang Y, Lu J, et al. A regeneration process-matching scaffold with appropriate dynamic mechanical properties and spatial adaptability for ligament reconstruction. Bioact Mater 2022;13:82–95. doi:10.1016/j.bioactmat.2021.11.001.

[20] Wang Z, Lee WJ, Koh BTH, Hong M, Wang W, Lim PN, et al. Functional regeneration of tendons using scaffolds with physical anisotropy engineered via microarchitectural manipulation. Sci Adv 2018;4:1–13. doi:10.1126/sciadv.aat4537.

[21] Benhardt HA, Cosgriff-Hernandez EM. The role of Mechanical loading in ligament tissue engineering. Tissue Eng - Part B Rev 2009;15:467–75. doi:10.1089/ten.teb.2008.0687.

[22] Voleti PB, Buckley MR, Soslowsky LJ. Tendon healing: Repair and regeneration. Annu Rev Biomed Eng 2012;14:47–71. doi:10.1146/annurev-bioeng-071811-150122.

[23] Vining KH, Mooney DJ. Mechanical forces direct stem cell behaviour in development and regeneration. Nat Rev Mol Cell Biol 2017;18:728–42. doi:10.1038/nrm.2017.108.

[24] Palanca M, Tozzi G, Cristofolini L. The use of digital image correlation in the biomechanical area: A review. Int Biomech 2016;3:1–21. doi:10.1080/23335432.2015.1117395.

[25] Luyckx T, Verstraete M, De Roo K, De Waele W, Bellemans J, Victor J. Digital image correlation as a tool for three-dimensional strain analysis in human tendon tissue. J Exp Orthop 2014;1:1–9. doi:10.1186/s40634-014-0007-8.

[26] Holak K, Kohut P, Ekiert M, Tomaszewski K, Uhl T. the Use of Digital Image Correlation in the Study of Achilles Tendon Strain Field. Mech Control 2015;34:19. doi:10.7494/mech.2015.34.1.19.

[27] Lozano PF, Scholze M, Babian C, Scheidt H, Vielmuth F, Waschke J, et al. Water-content related alterations in macro and micro scale tendon biomechanics. Sci Rep 2019;9:1–12. doi:10.1038/s41598-019-44306-z.

[28] Nagelli C V., Hooke A, Quirk N, De Padilla CL, Hewett TE, van Griensven M, et al. Mechanical and Strain Behaviour of Human Achilles Tendon During in Vitro Testing To Failure. Eur Cells Mater 2022;43:153–61. doi:10.22203/eCM.v043a12.

[29] Readioff R, Geraghty B, Comerford E, Elsheikh A. A full-field 3D digital image correlation and modelling technique to characterise anterior cruciate ligament mechanics ex vivo. Acta Biomater 2020;113:417–28. doi:10.1016/j.actbio.2020.07.003.

[30] Mallett KF, Arruda EM. Digital image correlation-aided mechanical characterization of the anteromedial and posterolateral bundles of the anterior cruciate ligament. Acta Biomater 2017;56:44–57. doi:10.1016/j.actbio.2017.03.045.

[31] Li X, Xie J, Lipner J, Yuan X, Thomopoulos S, Xia Y. Nanofiber scaffolds with gradations in mineral content for mimicking the tendon-to-bone insertion site. Nano Lett 2009;9:2763–8. doi:10.1021/nl901582f.

[32] Lipner J, Liu W, Liu Y, Boyle J, Genin GM, Xia Y, et al. The mechanics of PLGA nanofiber scaffolds with biomimetic gradients in mineral for tendon-to-bone repair. J Mech Behav Biomed Mater 2014;40:59–68. doi:10.1016/j.jmbbm.2014.08.002.

[33] Mubyana K, Koppes RA, Lee KL, Cooper JA, Corr DT. The influence of specimen thickness and alignment on the material and failure properties of electrospun polycaprolactone nanofiber mats. J Biomed Mater Res - Part A 2016;104:2794–800. doi:10.1002/jbm.a.35821.

[34] Caballero DE, Montini-Ballarin F, Gimenez JM, Biocca N, Rull N, Frontini P, et al. Reduced kinematic multiscale model for tissue engineering electrospun scaffolds. Mech Mater 2022;166:104214. doi:10.1016/j.mechmat.2022.104214.

[35] Bay BK. Texture correlation: A method for the measurement of detailed strain distributions within trabecular bone. J Orthop Res 1995;13:258–67. doi:10.1002/jor.1100130214.

[36] Dall’Ara E, Tozzi G. Digital volume correlation for the characterization of musculoskeletal tissues: Current challenges and future developments. Front Bioeng Biotechnol 2022;10:1–13. doi:10.3389/fbioe.2022.1010056.

[37] Marr N, Hopkinson M, Hibbert AP, Pitsillides AA, Thorpe CT. Bimodal Whole-Mount Imaging of Tendon Using Confocal Microscopy and X-ray Micro-Computed Tomography. Biol Proced Online 2020;22:1–14. doi:10.1186/s12575-020-00126-4.

[38] De Bournonville S, Vangrunderbeeck S, Kerckhofs G. Contrast-enhanced microCT for virtual 3D anatomical pathology of biological tissues: A literature review. Contrast Media Mol Imaging 2019;2019. doi:10.1155/2019/8617406.

[39] Marchiori G, Parrilli A, Sancisi N, Berni M, Conconi M, Luzi L, et al. Integration of micro-CT and uniaxial loading to analyse the evolution of 3D microstructure under increasing strain: Application to the Anterior Cruciate Ligament. Mater Today Proc 2019;7:501–7. doi:10.1016/j.matpr.2018.11.116.

[40] Tits A, Plougonven E, Blouin S, Hartmann MA, Kaux JF, Drion P, et al. Local anisotropy in mineralized fibrocartilage and subchondral bone beneath the tendon-bone interface. Sci Rep 2021;11. doi:10.1038/s41598-021-95917-4.

[41] Pierantoni M, Silva Barreto I, Hammerman M, Verhoeven L, Törnquist E, Novak V, et al. A quality optimization approach to image Achilles tendon microstructure by phase-contrast enhanced synchrotron micro-tomography. Sci Rep 2021;11:1–14. doi:10.1038/s41598-021-96589-w.

[42] Pierantoni M, Hammerman M, Silva I, Andersson L, Novak V, Isaksson H, et al. Journal of Structural Biology : X Heterotopic mineral deposits in intact rat Achilles tendons are characterized by a unique fiber-like structure. J Struct Biol X 2023;7:100087. doi:10.1016/j.yjsbx.2023.100087.

[43] Sartori J, Köhring S, Bruns S, Moosmann J, Hammel JU. Gaining Insight into the Deformation of Achilles Tendon Entheses in Mice. Adv Eng Mater 2021;23:1–11. doi:10.1002/adem.202100085.

[44] Kannus P. Structure of the tendon connective tissue. Scand J Med Sci Sport 2000;10:312–20. doi:10.1034/j.1600-0838.2000.010006312.x.

[45] Murphy W, Black J, Hastings G. Handbook of Biomaterial Properties. Second. Springer; 2016. doi:10.1007/978-1-4939-3305-1.

[46] Liu Z. Scale space approach to directional analysis of images. Appl Opt 1991;30:1369–73. doi:https://doi.org/10.1364/AO.30.001369.

[47] Schindelin J, Arganda-Carreras I, Frise E, Kaynig V, Longair M, Pietzsch T, et al. Fiji: an open-source platform for biological-image analysis. Nat Methods 2012;9:676–82. doi:10.1038/nmeth.2019.

[48] Sensini A, Cristofolini L, Focarete ML, Belcari J, Zucchelli A, Kao A, et al. High-resolution x-ray tomographic morphological characterisation of electrospun nanofibrous bundles for tendon and ligament regeneration and replacement. J Microsc 2018;272:196–206. doi:10.1111/jmi.12720.

[49] Gotti C, Sensini A, Fornaia G, Gualandi C, Zucchelli A, Focarete ML. Biomimetic Hierarchically Arranged Nanofibrous Structures Resembling the Architecture and the Passive Mechanical Properties of Skeletal Muscles : A Step Forward Toward Artificial Muscle. Front Bioeng Biotechnol 2020;8:767. doi:10.3389/fbioe.2020.00767.

[50] Dall’Ara E, Bodey AJ, Isaksson H, Tozzi G. A practical guide for in situ mechanical testing of musculoskeletal tissues using synchrotron tomography. J Mech Behav Biomed Mater 2022;133:105297. doi:10.1016/j.jmbbm.2022.105297.

[51] Maes A, Pestiaux C, Marino A, Balcaen T, Leyssens L, Vangrunderbeeck S, et al. Cryogenic contrast-enhanced microCT enables nondestructive 3D quantitative histopathology of soft biological tissues. Nat Commun 2022;13:1–14. doi:10.1038/s41467-022-34048-4.

[52] Stamati O, Andò E, Roubin E, Cailletaud R, Wiebicke M, Pinzon G, et al. spam: Software for Practical Analysis of Materials. J Open Source Softw 2020;5:2286. doi:10.21105/joss.02286.

[53] Dall’Ara E, Peña-Fernández M, Palanca M, Giorgi M, Cristofolini L, Tozzi G. Precision of digital volume correlation approaches for strain analysis in bone imaged with micro-computed tomography at different dimensional levels. Front Mater 2017;4. doi:10.3389/fmats.2017.00031.

[54] Johnson CR, Hansen CD. The Visualization Handbook. Elsevier; 2005.

[55] Hanson P, Aagaard P, Magnusson SP. Biomechanical properties of isolated fascicles of the Iliopsoas and Achilles tendons in African American and Caucasian men. Ann Anat 2012;194:457–60. doi:10.1016/j.aanat.2012.03.007.

[56] Butler DL, Kay MD, Stouffer DC. Comparison of Material Properties in Fascicle-Bone Units From Human Patellar Tendon and Knee Ligaments. J Biomech 1986;19:425–32. doi:https://doi.org/10.1016/0021-9290(86)90019-9.

[57] Prusa G, Bauer L, Santos I, Thorwächter C, Woiczinski M, Kistler M. Strain evaluation of axially loaded collateral ligaments: a comparison of digital image correlation and strain gauges. Biomed Eng Online 2023;22:1–14. doi:10.1186/s12938-023-01077-z.

[58] Kohut P, Holak K, Ekiert M, Młyniec A, Tomaszewski KA, Uhl T. The application of digital image correlation to investigate the heterogeneity of Achilles tendon deformation and determine its material parameters. J Theor Appl Mech 2021;59:43–52. doi:10.15632/jtam-pl/127903.

[59] Knaus KR, Blemker SS. 3D Models Reveal the Influence of Achilles Subtendon Twist on Strain and Energy Storage. Front Bioeng Biotechnol 2021;9:1–10. doi:10.3389/fbioe.2021.539135.

[60] Lee AH, Szczesny SE, Santare MH, Elliott DM. Investigating mechanisms of tendon damage by measuring multi-scale recovery following tensile loading. Acta Biomater 2017;57:363–72. doi:10.1016/j.actbio.2017.04.011.

