## Supplementary Material for "Full-field strain distribution in hierarchical electrospun nanofibrous Poly-L(lactic) acid and Collagen based scaffolds for tendon and ligament tissue regeneration: a multiscale study"

**File include:**

Tables: 8 (Table S1, Table S2, Table S3, Table S4, Table S5, Table S6, Table S7, Table S8)

**Table S1.** Mechanical properties of SB, RB and EHS.

| | $F_Y$ (N) | $F_F$ (N) | $\varepsilon_Y$ (%) | $\varepsilon_F$ (%) | $\sigma_Y$ (MPa) | $\sigma_F$ (MPa) | E (MPa) | $W_Y$ (J/mm$^3$) | $W_F$ (J/mm$^3$) |
|---|---|---|---|---|---|---|---|---|---|
| **Apparent** | | | | | | | | | |
| SB | 0.91±0.19 | 2.73±0.54 | SB=RB | SB=RB | SB=RB | SB=RB | SB=RB | SB=RB | SB=RB |
| RB | 1.82±0.37 | 5.46±1.08 | 3.5±0.11 | 67.8±13.4 | 3.67±0.80 | 11.0±2.3 | 108±30 | 0.007±0.001 | 0.54±0.20 |
| EHS | 8.41±1.14 | 16.9±2.3 | 8.97±3.01 | 46.9±5.3 | 1.61±0.31 | 3.24±0.77 | 20.7±5.90 | 0.008±0.004 | 0.11±0.03 |
| **Net** | | | | | | | | | |
| SB | - | - | - | - | SB=RB | SB=RB | SB=RB | SB=RB | SB=RB |
| RB | - | - | - | - | 11.7±0.58 | 35.0±3.0 | 341±30 | 0.021±0.001 | 1.72±0.50 |
| EHS | - | - | - | - | 10.2±1.55 | 20.6±3.87 | 133±37 | 0.05±0.03 | 0.69±0.17 |

**Table S2.** The significance of differences between SB, RB and EHS yield and failure forces assessed with an ANOVA 1 test followed by a Tukey post doc.

| | $F_Y$ (N) | $F_F$ (N) |
|---|---|---|
| SB vs RB | ns (p=0.1390) | * (p=0.0361) |
| SB vs EHS | **** (p<0.0001) | **** (p<0.0001) |
| RB vs EHS | **** (p<0.0001) | **** (p<0.0001) |

**Table S3.** The significance of differences between SB and RB (SB|RB) and EHS apparent and net mechanical properties assessed with an unpaired parametric t-test with Welch's correction.

| SB\|RB vs EHS | $\varepsilon_Y$ (%) | $\varepsilon_F$ (%) | $\sigma_Y$ (MPa) | $\sigma_F$ (MPa) | E (MPa) | $W_Y$ (J/mm$^3$) | $W_F$ (J/mm$^3$) |
|---|---|---|---|---|---|---|---|
| Apparent | * (p=0.0153) | * (p=0.0213) | ** (p=0.0026) | *** (p=0.0008) | ** (p=0.0025) | ns (p=0.5869) | ** (p=0.0082) |
| Net | - | - | ns (0.1074) | *** (p=0.0002) | **** (p<0.0001) | ns (p=0.0991) | ** (p=0.0062) |

**Table S4.** The significance of differences of SB and RB (SB|RB) and EHS apparent and net mechanical properties assessed with a ratio paired parametric t-test.

| | $\sigma_Y$ (MPa) | $\sigma_F$ (MPa) | E (MPa) | $W_Y$ (J/mm$^3$) | $W_F$ (J/mm$^3$) |
|---|---|---|---|---|---|
| **SB\|RB** | | | | | |
| App. vs Net | **** (p<0.0001) | **** (p<0.0001) | **** (p<0.0001) | *** (p=0.0002) | **** (p<0.0001) |
| **EHS** | | | | | |
| App. vs Net | **** (p<0.0001) | **** (p<0.0001) | **** (p<0.0001) | **** (p<0.0001) | **** (p<0.0001) |

**Table S5.** Axial and transversal strain of EHS membranes.

| Strain Values (%) | 0% | 1.5% | 3% | 5% | 7% | $\varepsilon_Y$ | $\varepsilon_F$ |
|---|---|---|---|---|---|---|---|
| **Membrane** Axial Strain (%) | 0 | 7.24±3.11 | 9.44±2.86 | 11.59±2.95 | 13.75±2.58 | 16.08±2.79 | 21.67±3.67 |
| Transversal Strain (%) | 0 | -4.45±1.75 | -6.09±1.76 | -8.70±0.82 | -11.14±1.06 | -8.18±1.06 | -13.58±0.89 |

**Table S6.** MicroCT mechanical and morphological properties of SB and EHS.

| | CT.F (N) | CT.Cr.Ar (mm$^2$) | CT.σ (MPa) | Po/Tot.Po (%) | Int.Po (%) | τ | Θ (°) |
|---|---|---|---|---|---|---|---|
| **SB** | | | | | | | |
| **Strain Step** | | | | | | | |
| 2% | 0.46±0.01 | 0.170±0.043 | 3.50±1.95 | 3.2±3.32 | - | 1.036±0.003 | 2.98±1.17 |
| 3% | 0.98±0.20 | 0.168±0.044 | 6.27±2.83 | 2.7±2.95 | - | 1.057±0.003 | 2.82±0.99 |
| 4% | 1.12±0.40 | 0.167±0.045 | 7.39±4.56 | 2.4±2.75 | - | 1.054±0.007 | 2.54±0.69 |
| 5% | 1.25±0.47 | 0.166±0.045 | 8.03±4.11 | 2.0±2.38 | - | 1.053±0.003 | 2.81±0.92 |
| 7% | 1.39±0.50 | 0.165±0.044 | 8.85±4.09 | 1.5±1.73 | - | 1.056±0.003 | 2.86±0.96 |
| **EHS** | | | | | | | |
| **Strain Step** | | | | | | | |
| 0% | 0.45±0.00 | 3.12±0.28 | 0.14±0.01 | 11.6±10.9 | 14.7±11.4 | 1.095±0.003 | 6.79±1.49 |
| 1.5% | 1.87±1.72 | 2.96±0.43 | 0.6±0.48 | 13.7±11.0 | 16.2±10.7 | 1.094±0.004 | 6.68±1.58 |
| 3% | 3.14±1.38 | 2.75±0.44 | 1.13±0.40 | 17.1±7.7 | 19.2±7.1 | 1.094±0.004 | 6.54±1.75 |
| 5% | 5.18±1.10 | 2.57±0.54 | 2.07±0.52 | 19.6±6.7 | 20.1±6.9 | 1.094±0.004 | 6.31±1.87 |
| 7% | 6.57±0.24 | 2.40±0.67 | 2.90±0.90 | 21.9±7.2 | 21.2±6.0 | 1.093±0.004 | 6.12±1.98 |

**Table S7.** Mean ± SD of axial displacement, $\varepsilon_{p1}$, $\varepsilon_{p3}$, $\varepsilon_D$ and maximum and minimum values for SB.

| **In Situ Strain Steps** | **2-3%** | **2-4%** | **2-5%** | **2-7%** |
|---|---|---|---|---|
| **SB_1** | | | | |
| Disp (mm) | 0.12±0.05 | 0.19±0.06 | 0.30±0.10 | 0.49±0.16 |
| $Disp_{max}$ (mm) | 0.23 | 0.29 | 0.45 | 0.72 |
| $\varepsilon_{p1}$ (%) | 1.72±1.02 | 2.08±1.22 | 3.1±1.62 | 4.98±2.4 |
| $\varepsilon_{p1max}$ (%) | 7.37 | 8.60 | 10.82 | 14.03 |
| $\varepsilon_{p3}$ (%) | -0.87±0.46 | -0.87±0.53 | -1.23±0.72 | -1.8±1.04 |
| $\varepsilon_{p3min}$ (%) | -3.2 | -4.27 | -4.4 | -6.11 |
| $\varepsilon_D$ (%) | 0.75±1.08 | 1.03±1.36 | 1.62±1.82 | 2.64±2.3 |
| $\varepsilon_{Dmax}$ (%) | 6.53 | 7.86 | 9.61 | 13.44 |
| **SB_2** | | | | |
| Disp (mm) | 0.07±0.04 | 0.13±0.08 | 0.18±0.13 | 0.29±0.19 |
| $Disp_{max}$ (mm) | 0.14 | 0.27 | 0.39 | 0.61 |
| $\varepsilon_{p1}$ (%) | 1.86±2.91 | 2.34±1.78 | 3.23±2.51 | 5.21±3.23 |
| $\varepsilon_{p1max}$ (%) | 8.93 | 9.97 | 15.49 | 21.72 |
| $\varepsilon_{p3}$ (%) | -1.31±1.66 | -1.23±0.76 | -1.38±0.61 | -1.86±1.01 |
| $\varepsilon_{p3min}$ (%) | -5.07 | -6.94 | -6.71 | -7.67 |
| $\varepsilon_D$ (%) | 0.24±3.64 | 0.71±2.16 | 2.6±3.27 | 1.33±2.97 |
| $\varepsilon_{Dmax}$ (%) | 5.28 | 6.92 | 12.52 | 18.23 |
| **SB_3** | | | | |
| Disp (mm) | 0.06±0.03 | 0.11±0.05 | 0.15±0.06 | 0.27±0.08 |
| $Disp_{max}$ (mm) | 0.1 | 0.02 | 0.22 | 0.48 |
| $\varepsilon_{p1}$ (%) | 1.29±0.79 | 2.17±1.2 | 2.83±1.23 | 4.15±1.34 |
| $\varepsilon_{p1max}$ (%) | 4.44 | 6.73 | 7.28 | 7.97 |
| $\varepsilon_{p3}$ (%) | -0.73±0.43 | -0.94±0.45 | -1.46±0.65 | -1.89±0.74 |
| $\varepsilon_{p3min}$ (%) | -4.56 | -2.29 | -4.61 | -4.55 |
| $\varepsilon_D$ (%) | 0.42±1.12 | 1.03±1.28 | 1.22±1.46 | 1.66±2.1 |
| $\varepsilon_{Dmax}$ (%) | 3.59 | 5.97 | 7.15 | 10.68 |

**Table S8.** Mean ± SD of axial displacement, $\varepsilon_{p1}$, $\varepsilon_{p3}$, $\varepsilon_D$ and maximum and minimum values for EHS.

| In Situ Strain Steps | 0-1.5% | 0-3% | 0-5% | 0-7% |
|---|---|---|---|---|
| **EHS_1** | | | | |
| Disp (mm) | 0.07±0.03 | 0.12±0.05 | 0.22±0.05 | 0.38±0.08 |
| $Disp_{max}$ (mm) | 0.13 | 0.23 | 0.35 | 0.46 |
| $\varepsilon_{p1}$ (%) | 3.52±2.76 | 3.88±2.77 | 6.03±4.33 | 6.98±4.67 |
| $\varepsilon_{p1max}$ (%) | 13.43 | 20.89 | 30.66 | 36.49 |
| $\varepsilon_{p3}$ (%) | -3.02±2.22 | -3.85±2.17 | -6.4±3.82 | -6.36±3.74 |
| $\varepsilon_{p3min}$ (%) | -14.49 | -16.45 | -26.85 | -25.03 |
| $\varepsilon_D$ (%) | 4.75±2.21 | 5.65±2.56 | 9.1±4.38 | 9.72±4.12 |
| $\varepsilon_{Dmax}$ (%) | 16.36 | 19.01 | 36.35 | 33.85 |
| **EHS_2** | | | | |
| Disp (mm) | 0.09±0.02 | 0.18±0.04 | 0.30±0.07 | 0.44±0.12 |
| $Disp_{max}$ (mm) | 0.15 | 0.23 | 0.45 | 0.78 |
| $\varepsilon_{p1}$ (%) | 1.97±1.57 | 4.18±3.68 | 5.86±4.39 | 8.11±5.75 |
| $\varepsilon_{p1max}$ (%) | 12.42 | 29.27 | 36.14 | 41.12 |
| $\varepsilon_{p3}$ (%) | -1.93±1.52 | -3.62±2.58 | -4.57±2.85 | -6.27±3.59 |
| $\varepsilon_{p3min}$ (%) | -13.49 | -17.28 | -19.02 | -22.18 |
| $\varepsilon_D$ (%) | 2.85±2.02 | 5.64±3.91 | 7.51±4.2 | 10.3±5.57 |
| $\varepsilon_{Dmax}$ (%) | 18.52 | 30.11 | 32.27 | 37.05 |
| **EHS_3** | | | | |
| Disp (mm) | 0.03±0.01 | 0.09±0.02 | 0.19±0.04 | 0.33±0.09 |
| $Disp_{max}$ (mm) | 0.07 | 0.19 | 0.27 | 0.50 |
| $\varepsilon_{p1}$ (%) | 1.08±0.99 | 1.97±1.89 | 3.56±2.96 | 6.69±6.46 |
| $\varepsilon_{p1max}$ (%) | 7.09 | 18.05 | 24.81 | 42.91 |
| $\varepsilon_{p3}$ (%) | -2.27±1.57 | -4.83±3.28 | -7.83±4.43 | -12.41±7.18 |
| $\varepsilon_{p3min}$ (%) | -11.19 | -21.76 | -28.83 | -33.78 |
| $\varepsilon_D$ (%) | 2.45±1.39 | 5.05±3.01 | 8.52±4.13 | 12.21±6.93 |
| $\varepsilon_{Dmax}$ (%) | 10.79 | 19.28 | 28.59 | 42.42 |
